# A molecular cell atlas of endocrine signalling in human neural organoids

**DOI:** 10.1101/2025.08.14.669814

**Authors:** Gaja Matassa, Marco Tullio Rigoli, Davide Castaldi, Alessia Valenti, Manuel Lessi, Nicolò Caporale, Sarah Stucchi, Benedetta Muda, Riccardo Nagni, Lisa Mainardi, Alessandro Melon, Amaia Tintori, Davide Bulgheresi, Michal Kubacki, Sebastiano Trattaro, Sara Evangelista, Pim Leonards, Human Cell Atlas Organoid Biological Network, Carlo Emanuele Villa, Cristina Cheroni, Giuseppe Testa

## Abstract

Hormonal signalling shapes the development of the human brain and its disruption is implicated in various neuropsychiatric conditions. However, a comprehensive and mechanistic understanding of how hormonal pathways orchestrate human neurodevelopment remains elusive. Here we present a multi-scale high resolution atlas of endocrine signalling in human neural organoids through systematic perturbations with agonists and inhibitors of seven key hormonal pathways: androgen (AND), estrogen (EST), glucocorticoid (GC), thyroid (THY), retinoic acid (RA), liver X (LX), and aryl hydrocarbon (AH). By integrating bulk and single-cell transcriptomics, high-throughput imaging and targeted steroidomics, we mapped the molecular and cellular consequences of their physiologically relevant perturbations. Retinoic acid exerted the most profound effect, promoting neuronal differentiation and maturation, consistent with its established role as a patterning factor. Our analysis further benchmarked neural organoids for in vitro endocrinology and neurotoxicology by confirming previously reported *in vivo* effects, such as induction of mTOR signalling by AND, alteration of disease relevant genes by GC and enhanced differentiation by TH. Furthermore, we observed that LX activation upregulates genes involved in cholesterol metabolism while AH inhibition promotes neuronal differentiation. We next uncovered extensive crosstalks between these endocrine pathways, as in the paradigmatic convergence induced by AND agonist and inhibitors of GC, TH, and LX, affecting genes related to protein folding and metabolic regulation, as also highlighted by weighted gene co-expression network analysis. Single-cell analyses pinpointed cell-type-specific responses to hormonal challenges, such as the caudalization of progenitors and neurons upon RA activation and the depletion of specific neurodevelopmental states upon AH activation. Finally, we dissected the cytoarchitectural and morphometric impact of hormonal perturbations and demonstrated that neural organoids possess active steroidogenic pathways that are functionally modulated by the tested compounds. This atlas provides a systematic quantification of the hormonal impact on human neurodevelopment, enabling the investigation of uncharted aspects in the developmental origins of neuropsychiatric traits. Through the empowering architecture of its knowledge base for iterative adoption by the community, this resource will thus be key to probe how environmental factors and genetic endocrine vulnerabilities contribute to neurodevelopmental outcomes, as well as to train advanced generative models for improving their predictive power on gene environment interactions in human neurodevelopment.

## Main

Human neurodevelopment is orchestrated by a complex interplay of genetic programs and environmental cues ^1,2^, with endocrine signalling playing a pivotal role. Hormons impact a wide array of key processes, along the proliferation, differentiation, migration and maturation of the neural lineage. Acting through nuclear or membrane-associated receptors, hormones thus sculpt the developmental antecedents of the neural circuits that underpin cognition, emotions and sociability throughout the lifespan ^3–5^. Disruptions in the finely tuned hormonal milieu of foetal neurodevelopment, whether due to genetic mutations or exposure to endocrine-disrupting chemicals (EDCs), can have profound and long-lasting consequences on neurodevelopmental trajectories and contribute to the pathogenesis of neuropsychiatric conditions, as we previously showed ^6^.

Despite decades of research illuminating the roles of hormones for human health, primarily through animal studies and clinical observations of endocrine disorders, a comprehensive, mechanistic understanding of how diverse hormonal pathways orchestrate the complexities of human brain development remains unknown ^7^. Our current knowledge is fragmented, hampered by the ethical impossibility of direct experimental perturbations in humans, practical challenges in simultaneously and non-invasively measuring dynamic hormonal and neural changes, and significant limitations in translating findings from animal models due to species-specific differences.

Recent advances in single-cell omics and human neural organoid technologies have provided unprecedented opportunities to dissect the molecular and cellular dynamics of human neurodevelopment ^8–11^. Recapitulating key and human-specific aspects of early brain development, neural organoids offer a uniquely powerful platform to investigate the effects of hormones on neurodevelopmental processes and to identify the underlying regulatory networks. This has recently allowed to uncover key insights about the molecular impact of glucocorticoid ^12–14^, androgen ^15^, and thyroid ^6,16^ signalling on human neurodevelopment. Moreover, recent innovations in single-cell multiplexing approaches ^10,17,18^, allowed to screen several signalling molecules, including hormones, to study their influence on the differentiation of pluripotent stem cells into various neural cell types within organoid models, and how combinations of morphogens can drive regional and cell fate specification ^19–23^, as well as to generate atlases of hormonal signalling across tissues ^24,25^. Importantly, these approaches are unlocking the possibility to train generative computational models to perform increasingly reliable virtual screenings ^26–28^.

However, there is a lack of a comprehensive and systematic study on the impact of endocrine signalling during human neurodevelopment, covering also understudied pathways beyond the role of estrogen, androgen, thyroid and steroidogenesis (EATS) ^29,30^, and investigating the crosstalks among them.

Here we present a comprehensive atlas of endocrine signalling during neurodevelopment, resulting from chronic exposure of human neural organoids to agonists and inhibitors of seven different endocrine pathways. By combining bulk and single-cell transcriptomics, targeted steroidomics and high-throughput imaging, we uncovered key aspects of hormonal roles in human neurodevelopment, identifying hormone-responsive gene networks, elucidating the impact of specific hormones on neurodevelopmental cell types and states and defining their crosstalks. This represents a foundational resource for investigating the molecular and cellular underpinnings of neuropsychiatric traits, the role of environmental factors, in particular endocrine disruptive chemicals ^31^, and of genetic endocrine liabilities ^32^ in neurodevelopmental pathogenesis, and more broadly the impact of environmental factors on human health, in line with the need to integrate exposomics into biomedicine ^33–37^.

### Hormonal perturbations modulate the transcriptomic landscape of neural organoids

To unravel the impact of hormonal modulation on neurodevelopmental processes, we chronically exposed neural organoids differentiated from two genetically validated, control human induced pluripotent stem cell (hiPSC) lines (CTL08, male and CTL04, female) to agonists and inhibitors of seven hormonal pathways that play an important role in regulating human brain development: androgen (AND), estrogen (EST), glucocorticoid (GC), thyroid (THY), retinoic acid (RA), liver X (LX), and aryl hydrocarbon (AH), along with negative controls (including vehicle and/or untreated controls depending on the assay), for a total of 32 experimental conditions, each with at least three replicates for each of the molecular and cellular assays profiled (Fig. 1A, nomenclature section in the methods).

**Figure 1.**
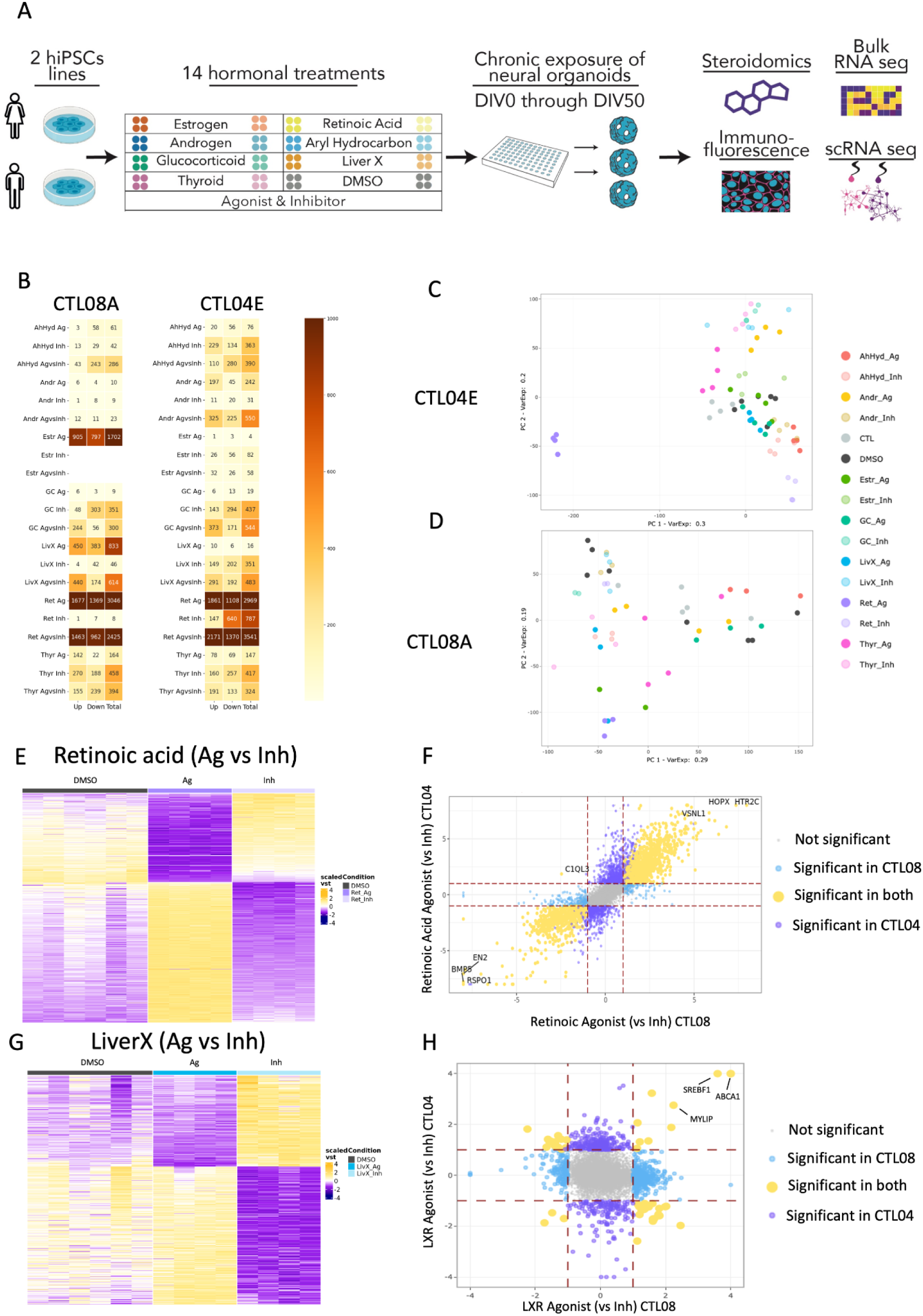
Experimental design and overall impact of hormonal perturbation on neural organoids bulk transcriptome. **A.** Experimental design summarising the entire study. **B.** Heatmaps showing the number of differentially expressed genes (DEGs) in the male (CTL08A) and female (CTL04E) cell lines. DEGs were identified from pairwise comparisons between each treatment, agonist (Ag) or inhibitor (Inh), and the DMSO control, as well as between each agonist and its corresponding inhibitor (Ag vs Inh). Genes were considered differentially expressed based on an adjusted p-value (FDR) < 0.01 and |log2 fold-change| > 1. The heatmaps indicate the number of upregulated (Up), downregulated (Down), and total DEGs for each comparison. **C-D.** Principal component analysis (PCA) of gene expression in exposed organoids from the female (CTL04E) and male (CTL08A) lines respectively. PC1 and PC2 are depicted. Samples are colored by treatment to highlight clustering patterns. **E-G.** Heatmap showing VST-normalized gene expression values (scaled by row) for DEGs from the RA Ag vs Inh comparison (E) or LX Ag vs Inh comparison (G) for CTL04E. DMSO-treated samples are included for reference. **F-H.** Comparison of DEGs identified in the RA Ag vs Inh condition (F) or LX Ag vs Inh between female and male lines. Light blue dots represent DEGs specific to the male line, violet to the female line, and yellow indicates shared DEGs, regardless of whether regulated in the same or opposite direction. The axes entail the log2 fold-changes.

Selection of the hormonal pathways was performed by verifying the expression of hormone receptors and key genes involved in hormones cellular uptake/prohormone activation in our neural organoid model that we previously characterized ^10,11^, and comparing it to the in-vivo counterpart (cortical samples from the Brainspan Atlas of the Developing Human Brain in bulk ^38^, and a single cell transcriptomic atlas of human neocortical development during mid-gestation ^39^) (Extended Data Fig. 1 and 2). Concentrations for the exposure experiments were chosen based on evidence of *bona fide* activation (for agonists) or inhibition (for inhibitors) of each hormonal pathway (Table 1). Organoids were exposed starting at induction of the neuroectodermal lineage (day in vitro, DIV 0) through DIV 50 (Fig. 1A). This timepoint is highly correlated with the first trimester of human fetal brain development ^9–11,40^, a critical period during which the developing brain is dependent on maternal hormonal cues as the fetal endocrine system is still largely immature ^7,41^. Given the inherent impossibility of recapitulating *in vitro* how the oscillatory patterns of systemic hormones translate into functional levels within individual cells, our design investigates, for each pathway, the full excursion between bona fide activation and inhibition. This interval is meant to capture, by definition, both physiological and physiopathological dose patterns (such as those arguably at play in chronic states of elevated or reduced activity in endocrine disorders), thereby serving as reference to study the full range of medically relevant hormonal effects on neurodevelopmental processes.

**Table 1.** Compounds utilized in this study and their corresponding concentrations.

Bulk transcriptomics profiling revealed for all tested compounds a significant albeit expectedly variable degree of modulation of the organoids’ transcriptome. To quantitatively characterize the molecular impact of endocrine perturbations, we systematically performed for each pathway i) supervised plotting and analysis of specific gene signatures, mainly related to neurodevelopmental processes and hormonal pathways; ii) differential expression analysis (testing both agonists and inhibitors vs DMSO to study the specific effects of each compound, as well as testing agonists vs inhibitors to capture the impact of the full endocrine axes perturbations); iii) functional enrichment analysis of the differentially expressed genes (DEGs) leveraging multiple curated knowledge bases and overlap analysis with disease-relevant databases; and iv) weighted gene co-expression network analysis (WGCNA) to identify convergent and divergent transcriptional programs across different endocrine pathways in an unsupervised manner.

We systematized this extensive dataset into a knowledge-base architecture (see methods) that allowed us to extract the salient findings while directly positioning them into a public, living atlas-scale resource to ground an iterative process of continuous refinement by the scientific community. Specifically, for each of the seven hormonal pathways, "report cards" available at [[GitHub report cards](https://github.com/GiuseppeTestaLab/noha/tree/main/1_bulkRNASeq/7_DataMining/Schemes)] distill the main observations drawn by our analyses for an entry level assessment, followed by a comprehensive multi-layered annotation that, as a key innovation, guides in depth, user-friendly exploration of our data within the context of the current body of scientific knowledge (dedicated pages at [[GitHub mining index](https://github.com/GiuseppeTestaLab/noha/tree/main/1_bulkRNASeq/7_DataMining/Schemes)]). Finally, given the higher variability across replicates of the same exposure condition observed for the organoids from the male line, our analysis focused on the female line first, which then served as a reference for a comparative interpretation of the data from the male line samples.

Among all perturbations, RA agonist was the compound that displayed the greatest transcriptional impact in both lines, as shown by the overall number of DEGs (Fig. 1B) and the clear segregation of this condition in the Principal Component Analysis (PCA) plot (Fig. 1C, D). This trend was largely shared between the two lines (Fig. 1F). Exposure to other compounds showed instead more line-specific effects, especially for androgen, estrogen, liver X, and aryl hydrocarbon modulators (Fig. 1H, Extended Data Fig. 3 B,D,L).

In the following paragraph, we highlight the results of transcriptomic analyses integrating observations from Fig. 1, Extended Data Fig. 3 and 4, and [[GitHub mining index](https://github.com/GiuseppeTestaLab/noha/blob/main/1_bulkRNASeq/7_DataMining/0.Index.md)]. These results mainly focus on the agonists vs inhibitors differential expression analysis, defined hereafter as Ag vs Inh.

In line with previous evidence regarding the pivotal role of RA signalling in neuronal differentiation ^42–44^, the RA Ag vs Inh analysis highlighted DEGs that are enriched for cell cycle-related categories (in the case of downregulated genes), and neuronal maturity and outer radial glia markers (for upregulated ones), together pointing to enhanced neuro- and astrogenesis, which in our non-exposed model was previously shown to peak at later developmental stages (> DIV100) ^10^.

Focusing on the other pathways, we observed that AND Ag vs Inh manifested downregulation of cell cycle genes, and an enrichment of upregulated DEGs for categories related to protein folding (a class of genes altered by many hormonal modulators) and mTOR signalling; interestingly an enrichment of cell cycle related categories was also observed among the downregulated DEGs for EST Ag vs Inh. GC Ag vs Inh, as opposed to AND Ag vs Inh, down-regulated genes enriched for categories related to protein folding, as well as nucleosome assembly. THY Ag vs Inh resulted in downregulation of cell cycle genes and protein folding categories, and increased expression of neuronal and oligodendrocyte markers as well as genes belonging to synaptic categories. Of note, cholesterol biosynthesis-related categories were significantly enriched among the DEGs across most of the exposure conditions. Investigating pathways beyond EATS ^30^, such as LX (extensively studied for the liver, but also relevant in neurobiology because of its involvement in brain cholesterol homeostasis ^45^), and Aryl hydrocarbon (active in neural cell types but still understudied in the nervous systems ^46^), we observed that the perturbation of both had a strong transcriptional impact. In particular, both LX and AH Ag vs Inh altered cell cycle genes expression and, while lipid metabolic pathways were enriched among the DEGs upregulated in the LX Ag vs Inh comparison, neuronal signaling and neurotransmission related pathways were enriched among the DEGs downregulated in the AH Ag vs Inh.

Overall, we provide a quantitative analysis of how different hormonal pathways shape the transcriptomic landscape of neurodevelopmental processes at the tissue level, demonstrating the power of neural organoids in both recapitulating well established mechanisms and disclosing new insights on the neurodevelopmental effects of endocrine pathways.

### Hormonal crosstalk in neurodevelopment

Since hormones can bind multiple receptors and activate multiple downstream effectors ^47–49^, studying hormonal crosstalk is essential for deciphering the complex mechanisms that guide human neurodevelopment. Our transcriptomic results showed strong convergences on specific functional categories (cholesterol biosynthesis and protein folding among others) across most of the endocrine modulators tested. We thus systematically analysed the overlaps between the DEGs of all agonists and inhibitors tested.

First, we found that in the female line significant overlaps were higher between DEGs altered by inhibitors than between those altered by agonists, with an opposite scenario for the male line (Fig. 2A, Extended Data Fig. 5B-E). Second, a strong convergence was observed in the female line between TH, GC and LX inhibitors, for both up- and down-regulated DEGs (Extended Data Fig. 5D), as also shown by the robust patterns in the heatmaps of Extended Data Fig. 5A. The significant overlap between TH and GC inhibitors down-regulated DEGs was also consistent in the male line (Fig. 2B, Extended Data Fig. 5E), and in line with a previous study in mice ^50^. Interestingly, in the female line AND Ag vs Inh induced the up-regulation of genes that significantly overlap with genes downregulated by all three pathways mentioned before (Fig. 2A). Finally, in both lines, DEGs altered by the RA agonist showed significant overlaps with different hormonal perturbations, highlighting a convergence with TH and AND in the female line and LX and EST in the male line (Extended Data Fig. 5B,C). Notably, for both lines, genes upregulated in the RA Ag vs Inh analysis significantly overlap with genes downregulated in the AH Ag vs Inh analysis, in line with the enrichment of neural differentiation-related genes upon RA activation and AH inhibition observed above (Fig. 2A,B).

**Figure 2.**
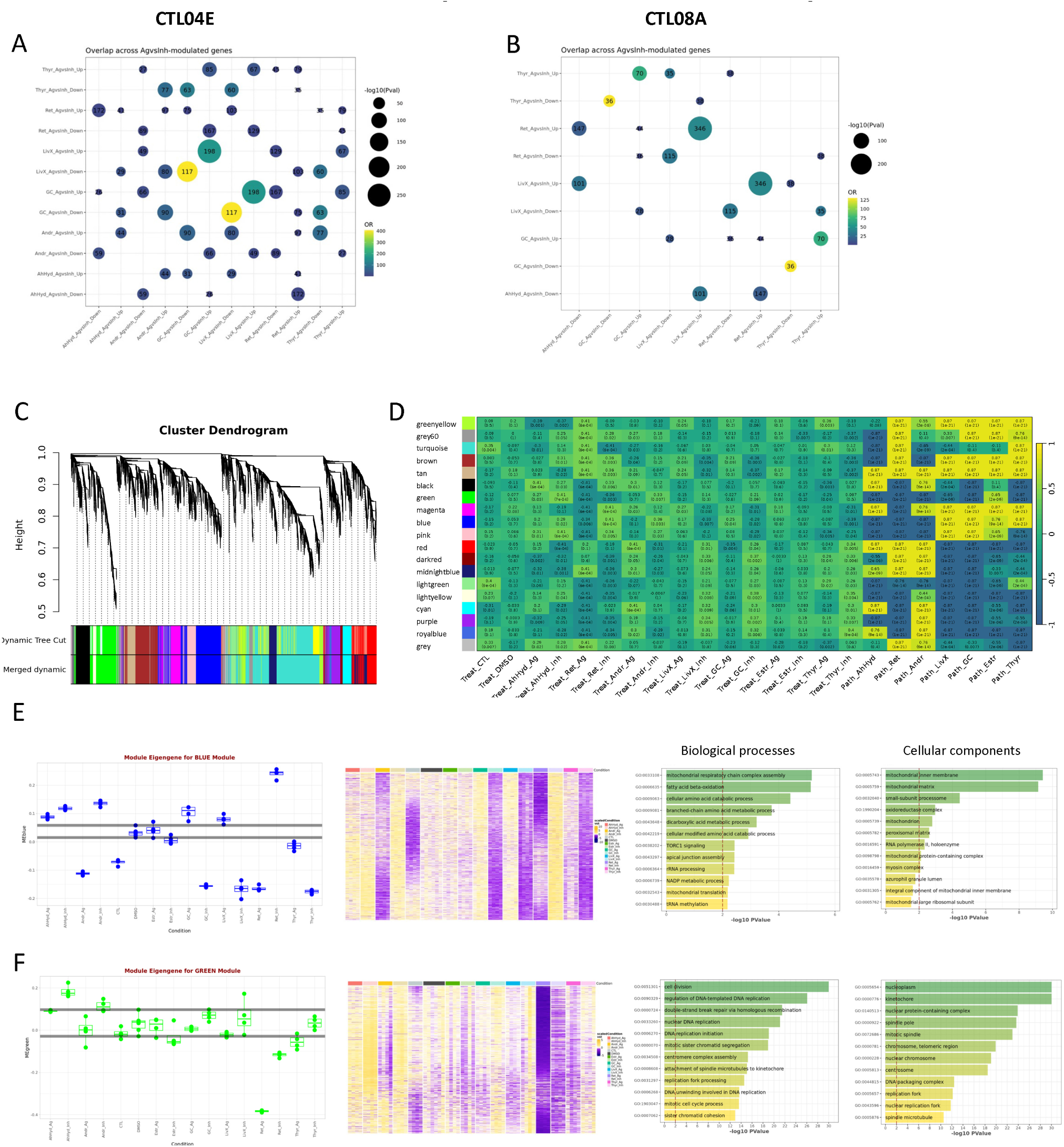
Hormonal crosstalk in neurodevelopment based on DEGs overlaps and WGCNA. **A-B**. Bubble plots showing DEGs overlap (Ag vs Inh) in the female and male line, respectively. Numbers represent shared genes; dot colour is assigned according to odds ratio (OR) values and dot size according to p-value as shown by each legend. Significant overlaps were selected according to the following thresholds: odds ratio (OR) > 2, p-value < 0.01, number of overlapping genes > 25. Matrices are symmetrical. **C**. Gene dendrogram of co-expressed gene modules generated by WGCNA workflow on the Topological Overlap Dissimilarity matrix for CTL04E samples. The rows below the dendrogram depicts module assignment before and after module merging, for a final total of 18 modules. **D**. Heatmap showing Spearman correlations between each gene module and individual treatments or pathways. **E,F.** Analysis of the blue (I) and GREEN (L) modules: left, boxplot of module eigengene values (the first principal component summarizing expression of all genes in the module) across treatments; dark grey horizontal lines represent the most extreme DMSO samples, included for reference. Middle, heatmap of VST-normalized gene expression values (scaled by row) for module genes across all CTL04E samples. Right, bar plots of the top 12 significantly enriched Biological Process and Cellular Component GO terms for the genes of the module. The red dashed line marks the p-value threshold at 0.01 (−log10(p-value) = 2).

To define the patterns of gene co-expression and their modulation by hormone agonists or inhibitors in an integrative manner, we performed WGCNA on the female line organoids. We identified and analyzed 18 gene modules. Many of them were significantly associated with multiple hormonal treatments (Fig. 2G,H). The gene composition, modulation across treatment, and functional characterization of the modules are thoroughly described in a dedicated section of the repository [[GitHub link](https://github.com/GiuseppeTestaLab/noha/blob/main/1_bulkRNASeq/7_DataMining/4.WGCNA.md)]. Among the modules indicating cross-talks of several treatments, the green one (Fig. 2F), functionally associated with DNA replication and cell division, was down-regulated by retinoic agonist and (less strongly) retinoic antagonist, but up-regulated by Aryl Hydrocarbon Antagonist. The brown (Extended Data Fig. 5G) and blue modules (Fig. 2E), functionally related to metabolism terms such as mitochondrial respiration (for both), fatty acid oxidation and amino acid catabolism (for the blue module) and triglycerides biosynthesis (for the brown), showed a widespread modulation across treatments, and a divergent response to RA agonist (brown module up-regulated by both RA agonist and antagonist and down-regulated by AND agonist and GC, LX and TYR inhibitors; blue module strongly upregulated by the inhibition of RA and downregulated by the activation of AND, RA, and by the inhibition of GC, LX, TH).

Several additional modules, capturing convergent as well as divergent hormonal actions in specific pathways, were identified such as the red module (Extended Data Fig. 5M), enriched for categories involved in protein folding and unfolded protein response, upregulated by RA and AND agonist, LX, GC and TH inhibitors; the cyan (Extended Data Fig. 5F) and royal blue (Extended Data Fig. 5L) modules enriched for epigenetic regulation terms, together confirming the interesting convergence between AND agonist, and LX, GC and TH inhibitors. Finally, two modules captured transcriptional programs that are much more pathway-specific, pinpointing alterations related to RA agonist: the grey60 (Extended Data Fig. 5I) module enriched for patterning factors, including several homeobox genes, and the turquoise module (Extended Data Fig. 5H), characterized by genes associated with neuronal maturation, such as synapse function and regulation.

In summary, our analyses identified and characterised the complex and multifaceted crosstalk between endocrine pathways acting during human neurodevelopment.

### Cell-type specific impact of hormonal perturbations on neurodevelopmental cell-states and gene modules

Next we undertook an in depth multiplexed single-cell transcriptomic profiling of the same conditions described above to dissect the cell-type-specific impact of hormonal perturbations on neurodevelopment. We performed demultiplexing by cell assignment to sample-specific barcodes ^51^ and deconvolution of genetic identities with SCanSNP genetic consensus call, which we previously developed and benchmarked ^10^, to maximize the call reliability and to pre-filter doublets and low-quality cells. Individual pre-processing and filtering of each treatment condition resulted in a total of 157,197 high-quality cells. Notably, consistent differences between cell lines were observed, with low variability across exposure replicates (Extended Data Fig. 6A). Therefore, after an initial dimensionality reduction and integration ^52^ for exploratory analysis and annotation, downstream quantitative analyses were performed separately for each line, without integration, to preserve line-specific differences. Exploratory analysis of the subset of cells from the vehicle control condition (DMSO), confirmed that organoids from both lines recapitulated expected cell types (Extended Data Fig. 7A,B). We then annotated the main cell types (neural progenitors and neurons) on the entire dataset, by analysing the expression distribution of relevant markers and projecting the cells onto human fetal brain references (Extended Data Fig. 6A,B). We first observed that for both hiPSC lines, in line with the results from the bulk analyses, organoids treated with RA agonist had lower number of cycling cells and exposure to EST inhibitor and RA inhibitor increased the number of cycling cells (Extended Data Fig. 6C). We then proceeded to study the specific impact of hormonal perturbations on each cell type. Exploratory analysis of marker gene expression suggested that hormonal exposure biased towards diverse states of progenitors and neurons (Fig. 3). To quantitatively map these effects, we performed differential abundance analysis between agonists and inhibitors-treated cells for each hormonal pathway using Milo, a cluster-free statistical approach ^53^. We observed that some conditions were significantly depleted in specific cell states, such as AH activation for LHX9-positive pioneer-like neurons, which *in vivo* settle in the marginal zone during early stages of corticogenesis ^54,55^. On the other hand, RA agonist was significantly enriched for caudalised progenitors and neurons marked by HOXB9. Interestingly, AND activation induced enrichment for different states of neurons and progenitors only in the male line, in contrast to what had emerged from bulk transcriptome analysis with the female line more heavily impacted by AND (Fig. 3, Extended Data Fig. 8).

**Figure 3.**
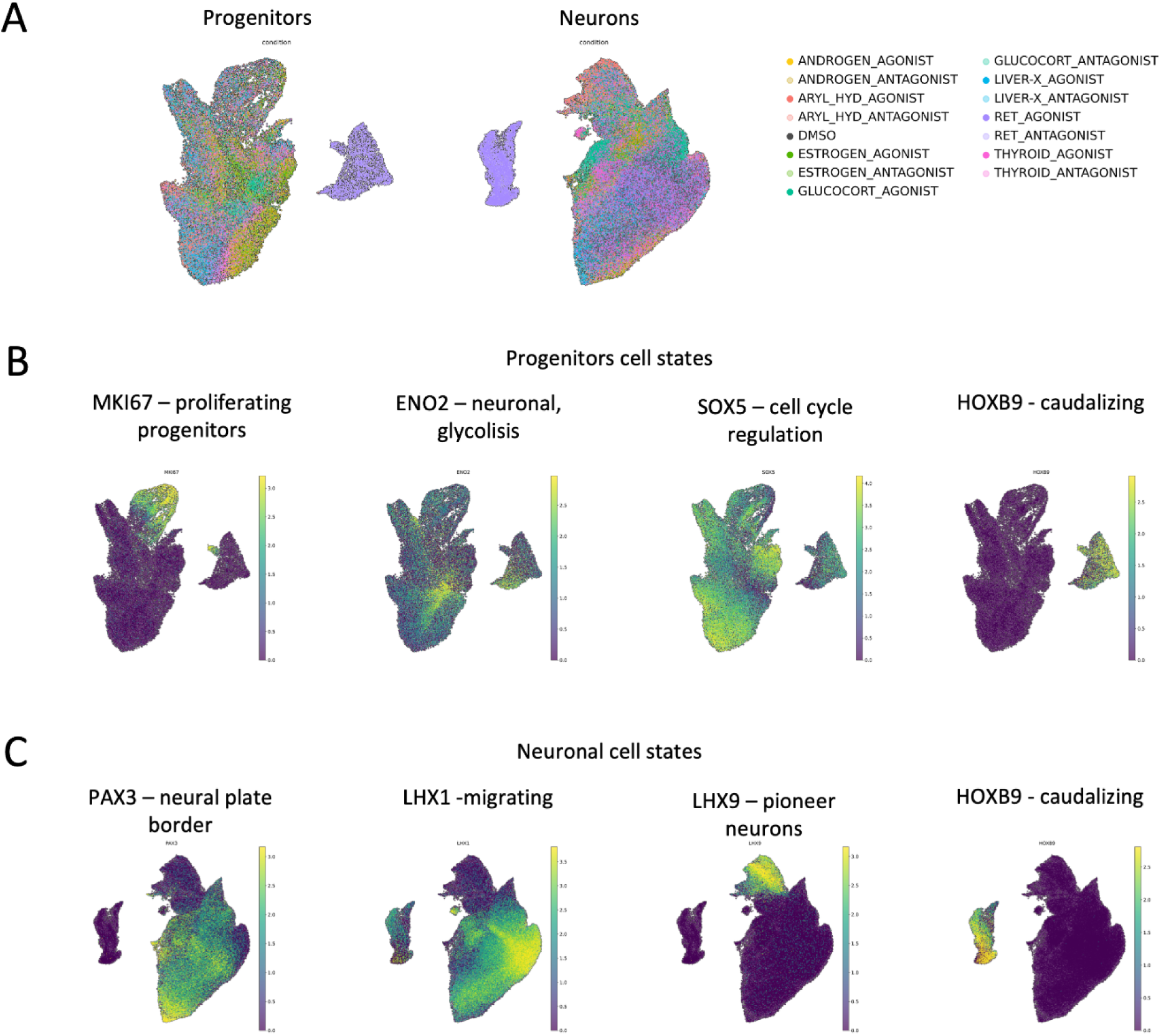
Single cell dissection of hormonal impact on cell states. **A.** Cell embeddings in UMAPs after preprocessing, filtering and annotation of progenitors and neurons. Each dot is a cell, colored by exposure condition. **B.** UMAPs of the progenitors coloured by expression of relevant markers. **C.** UMAPs of the neurons coloured by expression of relevant markers.

These results prompted us to investigate the impact of AND agonist on neural organoids from four different hiPSC lines (2 males and 2 females) in a new independent single cell experiment recapitulating the same experimental condition, chronically exposing organoids through 50 days to AND agonist. After verifying cell’s quality and the recapitulation of the expected cell types across all conditions and lines, we found no significant difference in this experiment across lines or across sexes upon AND agonist exposure (Extended Data Fig. 7 C,D, [[GitHub link](https://giuseppetestalab.github.io/noha/10.1_Androgen_substudy_DA.html)]).

Finally, pseudobulking cells (aggregating cells of the different replicates for each line, condition and cell type), we performed WGCNA to capture the most interesting gene modules of co-expression across exposure conditions, cell types and lines. We identified modules enriched for lipid metabolism, telomere regulation, and steroid hormone signalling pathway (orange-red and maroon from the neurons), as well as cellular metabolism (lemonchiffon from the progenitors), that were altered in several hormonal treatments in the same direction across the two lines (Fig 4). Moreover, confirming previous results, in particular the grey60 module from the bulk WGCNA, both the saddlebrown module in progenitors and the darkgrey module in neurons highlighted HOX genes related to patterning and regionalisation enriched for RA agonist treated cells (Fig 4). Detailed characterisation of all modules is available here [[GitHub link for progenitors](https://giuseppetestalab.github.io/noha/11.1_Hormones_substudy_LineMagnification_Progenitors_DA_divided.html), [GitHub link for neurons](https://giuseppetestalab.github.io/noha/11.2_Hormones_substudy_LineMagnification_Neurons_DA_divided.html)].

**Figure 4.**
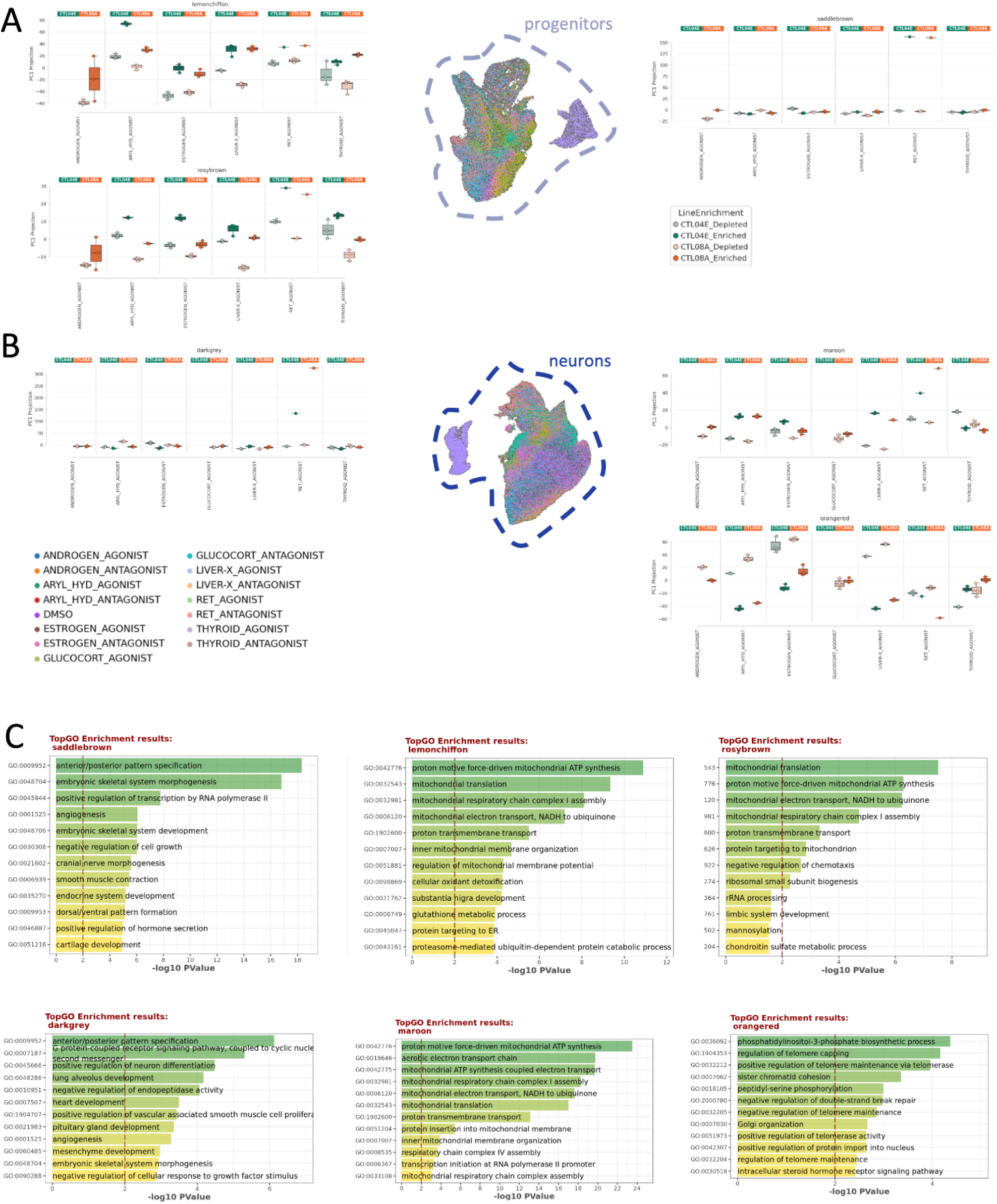
Cell-type specific impact of hormonal perturbations through co-expressed gene modules. **A**. Selected examples of gene modules modulated by several hormonal perturbations (lemonchiffon and rosybrown modules), or specifically by RA (saddlebrown module) from weighted gene co-expression network analysis (WGCNA) on progenitors of both cell lines. **B**. Selected examples of gene modules modulated by several hormonal perturbations (maroon and orangered modules), or specifically by RA (darkgrey module) from weighted gene co-expression network analysis (WGCNA) on neurons of both cell lines. **C**. Bar plots displaying the top 12 enriched Biological Process GO terms (ranked by p-value) for the genes of each WGCNA module shown in panel A and B. The red vertical dashed line indicates the significance threshold at p-value = 0.01 (−log10(p-value) = 2).

This single-cell characterization identified cell-type-specific effects of hormonal perturbations, identifying distinct cellular states and gene modules that are selectively and/or convergently modulated by different endocrine pathways during neurodevelopment.

### High-throughput imaging reveals the impact of hormonal perturbations on organoids cytoarchitecture and morphometrics

To elucidate the impact of hormonal perturbations on neurodevelopment on a complementary level, we performed high-throughput imaging on neural organoids from the same round of differentiation profiled for single cell transcriptomics. To minimise technical variability and to allow the parallel processing of a high number of exposure conditions, organoids fixed from each exposure condition were assembled into paraffin-embedded Tissue Micro Arrays (TMA). Upon immunofluorescence staining for two progenitor and proliferation markers (respectively, SOX2 and KI67), and two neuronal markers (MAP2 and TUJ1) (Table 2), images were acquired through digital slide scanning, and a semi-automated pipeline (including deep learning based nuclei/cell detection and segmentation, and multi-tiered manual quality checks to discard all cases where tissue quality was not preserved) was developed and applied [[GitHub link](https://github.com/GiuseppeTestaLab/noha/tree/main/3_ImageAnalysis/preprocessing)]. This approach enabled a quantitative assessment of 146 neural organoids, and ∼ 5 million nuclei (Fig. 5A, Extended Data Fig. 9A). We observed that both the percentage of progenitors and the organoid area covered by neuronal markers, normalized on the total number of nuclei, were higher in organoids treated with RA and, to a smaller extent, also for organoids treated with AND modulators, compared to controls (Extended Data Fig. 9D). This indicated an alteration of the cytoarchitecture of the organoids. Indeed, both conditions showed lower cellular density within the organoids (number of nuclei/organoid’s area) (Extended Data Fig. 9C). This represents an interesting finding that could only be captured as a cytoarchitectural phenotype by imaging, complementing the previous omics findings. In addition, we leveraged the segmentation of organoids’ nuclei to study the impact of hormonal exposure on nuclear morphological features. Morphometric feature extraction allowed us to perform principal component analysis and stratify the organoids of each exposure according to these axes of variation (Fig. 5B). RA activation confirmed to be the perturbation with the strongest impact, also on this layer of analysis, especially considering the first principal component, which is mainly driven by nuclear size-related features. Interestingly, higher nuclei size was observed for organoids exposed to RA agonist and, to a smaller extent, for AND modulators (Extended Data Fig. 9E), complementing the previous results on progenitors and neurons and on the organoids cytoarchitecture. Finally, several hormonal treatments, especially AND, AH, and LX modulations, had an impact on nuclear shape-related features, as highlighted by the second principal component (Fig. 5B).

**Figure 5.**
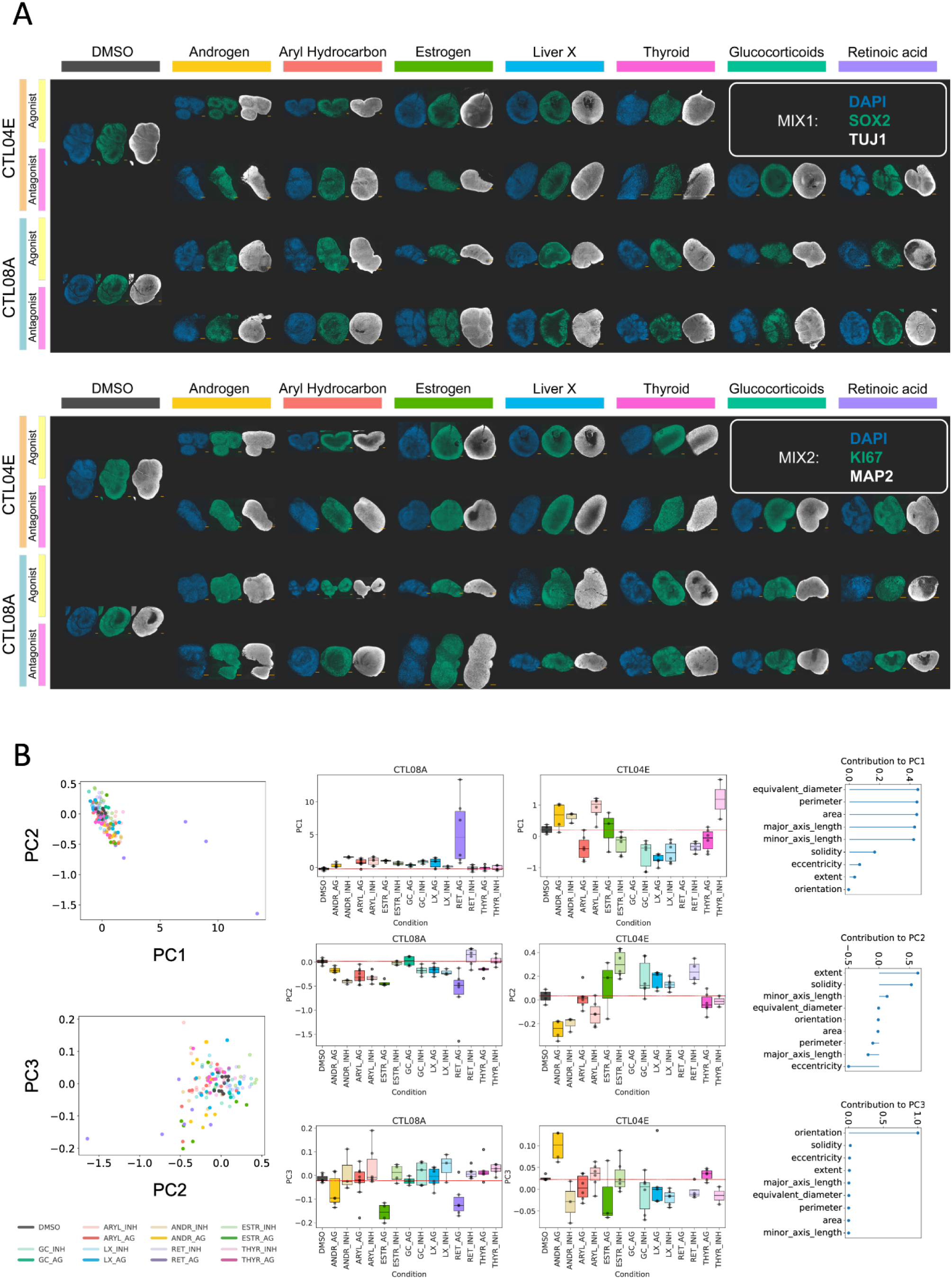
Hormonal impact on neural organoids cytoarchitecture and morphometrics. **A.** Representative images of immunostainings with two different panels of primary antibodies from each condition divided by line. In the bottom right corner, the orange scale bar depicts 200 µm. **B.** Principal component analysis performed on the morphological features of all nuclei and averaged per organoid. On the left side, scatterplots of the averaged values of PC1 and PC2 (top) and of PC2 and PC3 (bottom) colored by condition. In the middle, boxplots of average value of PC1 (top), PC2 (middle) and PC3 (bottom) per organoid across conditions and divided by line. Box whiskers show the first and third quartiles of the data and the black solid line the median. The red dotted line in each plot represents the median of the DMSO condition. On the right side, lollipop plots show the signed contributions of the covariates to PC1 (top), PC2 (middle) and PC3 (bottom).

**Table 2.** List of primary antibodies used to perform immunofluorescence stainings on neural organoids.

These findings provide evidence that hormonal signalling impacts neural organoids’ cell type composition and cytoarchitecture, as well as nuclear morphotypes.

### Hormonal perturbations modulate steroid metabolic pathways in neural organoids

Steroidogenesis is the biochemical process by which steroid hormones are synthesised from cholesterol ^56^. Steroid local synthesis in neural cells is relevant for neurodevelopmental processes and can be influenced by the hormonal modulators we tested ^57,58^. For each exposure condition, we thus profiled steroid metabolites using a targeted approach to dissect the impact of hormonal perturbations on neural organoids steroidogenesis (Fig 6, Extended Data Fig 10, Table 3).

**Figure 6.**
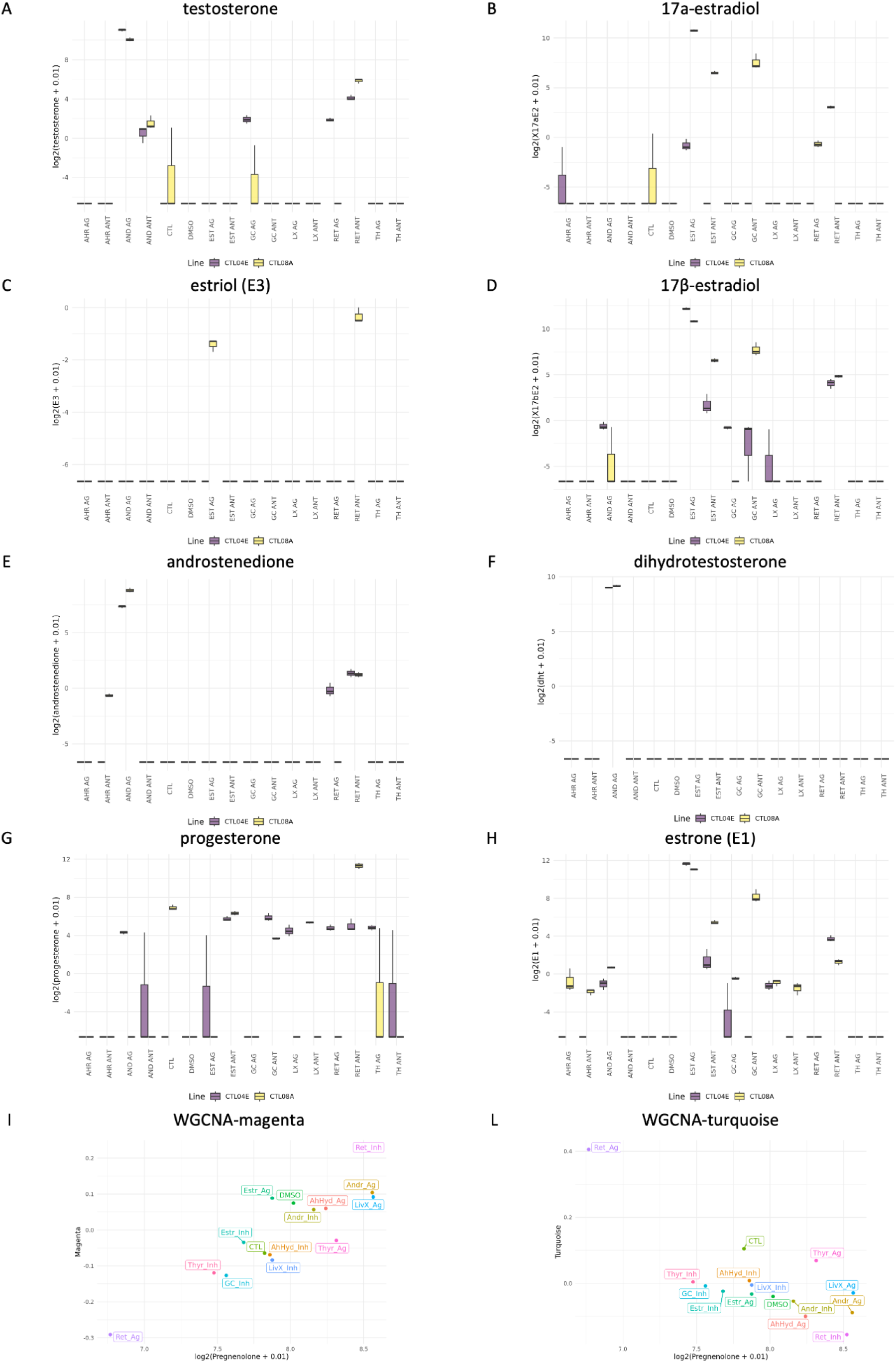
Steroidogenesis profiling in neural organoids upon endocrine perturbations. **A-H**. Boxplots of the log2 (value + 0.01) of the measured steroid metabolites in neural organoids from both hiPSC lines exposed to endocrine agonists and inhibitors. **I-L.** Correlation plot between the eigengene values of the magenta and turquoise modules identified by WGCNA, and the mean log2values (as calculated in A-H) of pregnenolone in each condition measured in neural organoids exposed to endocrine agonists and inhibitors.

**Table 3.** Multiple Reaction Monitoring (MRM) transition table and Source and Acquisition Parameters (SCIEX) for targeted steroids analysis.

We first confirmed that pregnenolone, the precursor in the biosynthesis of most of the steroid hormones, was high in all conditions, demonstrating active steroidogenesis (Extended Data Fig 10B). Second, organoids exposed to sex hormones (AND and EST agonists) showed much higher levels of testosterone and 17β-estradiol relative to the other exposure conditions (Fig 6A,D), demonstrating that organoids can effectively internalise exogenous hormones during development. Next, we observed that both lines endogenously produced testosterone upon exposure to RA and AND inhibitors, and by the female line upon RA and GC agonists (Fig 6A). While RA and GC alterations have previously been linked to steroidogenesis ^59–62^, our data represent the first evidence of testosterone production in response to AND inhibitor. Testosterone can be synthesized from androstenedione via 17β-hydroxysteroid dehydrogenases or from androstenediol via 3β-hydroxysteroid dehydrogenase. We observed an upregulation of androstenedione upon RA antagonist and AND agonist exposure, but not upon AND antagonist exposure (Fig 6E). This suggests that testosterone may increase through androstenediol upregulation under AND antagonism. Moreover, dihydrotestosterone (DHT), the active androgen synthesised from testosterone, was detectable only upon AND agonist (Fig 6F). This suggests that chronic AND overactivation is required to trigger the enzymatic machinery necessary for this conversion. As for estrogens, we observed that estrone was elevated upon EST agonist and to a lesser extent upon EST and RA inhibition, as well as AND and LX agonism (Fig 6H). Since estrone is derived from androstenedione via aromatase, its increase upon RA inhibition and AND agonism could be a consequence of an excess of androstenedione (Fig 6E), whereas the link with the LX pathway was not previously known. Both 17α-estradiol and 17β-estradiol were strongly upregulated in both lines upon estrogen activation. Of note, 17β-estradiol was also modestly increased following AND agonism (Fig 6B,D). In contrast, estriol was detected only upon EST agonism and RA inhibition in the male line (Fig 6C), indicating a possible line-specific ability to convert estrone and estradiol into this terminal estrogen metabolite.

Interestingly, among the measured metabolites, corticosterone and progesterone were the only ones present in the B27 supplement ([link](https://www.thermofisher.com/it/en/home/technical-resources/media-formulation.250.html)), which is part of the culture media and were thus constantly provided to the organoids. While corticosterone was indeed high in all exposure conditions (Extended Data Fig 10C), progesterone levels increased only following exposure to EST, GC, and RA antagonists (Fig 6G).

Finally, performing an analysis across scales, we examined the relationship between the WGCNA modules from the bulk transcriptomics and the levels of compounds measured by steroidomics. Among the four steroids which concentration was significantly correlated with some WGCNA modules, the most interesting were found for pregnenolone, positively correlated with the magenta module, which is enriched for extracellular matrix-related genes, and negatively correlated with the turquoise module, which is enriched for neuronal maturation-related genes (Fig 6I,L, [[GitHub link](https://www.thermofisher.com/it/en/home/technical-resources/media-formulation.250.html)]).

Together, these results demonstrate that neural organoids possess the molecular machinery required for steroid synthesis and metabolism and complement transcriptomic profiling to reveal functional hormonal crosstalk across different pathways.

## Discussion

With this study we introduce a comprehensive high resolution, multi-scale atlas of endocrine signalling in human neurodevelopment by systematically perturbing human neural organoids with agonists and antagonists of seven different hormonal pathways.

Our results showed RA signalling exerted the most profound molecular and cellular impact on neural organoids. This was evident across all scales, including i) the downregulation of cell-cycle genes in bulk transcriptomics and the reduced proportion of cycling cells upon single cell analysis, in line with a recent study adopting the neurosphere assay as an *in vitro* test method for developmental neurotoxicity evaluation ^5^; ii) the robust increase of neuronal and glial differentiation and maturation in bulk transcriptomics coupled with altered cytoarchitecture and increased neuronal marker expression observed from imaging; and iii) the strong caudalised signature of progenitors and neurons upon single cell analysis, in line with the results from our bulk transcriptomics and previous studies ^63,64^. These findings are consistent with the established pivotal role of RA in orchestrating neural patterning and differentiation, thereby demonstrating the sensitivity of this neural organoid model in detecting and validating RA regulatory effects from established in vivo systems ^22,42–44^.

Moreover, by comparing our findings with recent studies that investigated the role of specific hormonal pathways on human neurodevelopment through neural organoids modelling, we confirmed:

i. the induction of the mTOR signaling pathway upon AND activation, in line with ^15^;
ii. the impact of GC perturbation on neurodevelopmental disorders causative genes, in line with ^14^, and the FKBP5 relay, in line with post-traumatic stress disorder ^65^. Instead, our data did not replicate the previously observed effects of GC alteration on interneurons, possibly due to different organoid protocols and stages profiled ^14^;
iii. the increased differentiation induced by TH agonist, in line with ^6,16^, including for the oligodendrocyte lineages, consistent with the use of thyroid hormone for the differentiation of oligodendrocyte-enriched neural organoids ^66,67^.

Beyond EATS ^30^, our atlas illuminated the role of pathways that are less characterized in the context of human neurodevelopment. As expected from animal studies, activation of LX led to upregulation of genes central to cholesterol metabolism and transport, most notably APOE, in line with ^68^. Moreover, we discovered a convergent impact of LX inhibition (shared with AND agonist as well as GC and TH inhibitors) on protein folding and epigenetic regulation-related genes, thus providing novel evidence for an important developmental role of LX, and, more broadly, a previously unknown convergence across these four hormonal pathways on neurodevelopment. This complements the already known crosstalk between GC and AND, whose receptors share core DNA binding sites ^59^. Finally, inhibition of AH, whose receptor was previously defined as a xenobiotic sensor ^69^, led to an enrichment of neuronal differentiation genes in our organoids and indeed also showed a significant overlap with RA agonist DEGs. AH activation led instead to the depletion of a subpopulation of LHX9-expressing neurons from the single-cell analysis. These findings provide a potential mechanism to explain the neurodevelopmental defects observed in zebrafish following AH activation ^70^, and the epidemiological associations between exposure to potent AH agonists like dioxins and fetal alterations ^71^.

In line with a recent work investigating the impact of morphogens exposure on neural organoids that showed substantial interindividual and line-to-line variations in response to morphogens ^21^, our results also revealed a variable response of the two lines to hormonal perturbations. This was different across pathways and layers of analysis. For example the RA agonist produced very similar effects on the two lines across all the assays, whereas other compounds, such as AND, EST, LX, and AH modulators, showed line-specific effects. Interestingly, our results are coherent with the observation from ^21^ that showed inter-line variability to be higher in WNT-response than in SHH-response, indicating complex gene-environment interactions when translating environmental cues into cellular developmental programs. While our results could also be the consequence of interesting genetic sex-specific mechanisms ^72,73^, future neural organoid studies on a higher number of lines are needed to address these aspects, as underlined by the absence of significant differences when we tested AND agonist across multiple lines.

In conclusion, this atlas provides an unprecedented multi-scale quantitative reference of endocrine signalling in human neurodevelopment. We have confirmed known hormonal roles, uncovered novel neurodevelopmental functions for understudied endocrine pathways, and mapped the extensive crosstalk between them. Together, the density of this dataset enabled us to define the hallmarks of endocrine signalling in human neural organoids (Fig. 7), establishing a foundational resource for investigating the developmental landscape of neuropsychiatric traits and for analysing how environmental exposures shape human neurodevelopment ^25,33,35–37^. Importantly, this also represents a valuable resource for clinicians to investigate the molecular mechanisms underlying neuropsychiatric complications observed in endocrine disorders, where the developing human brain is exposed to hormonal imbalances, such as hyper- and hypo-thyroidism, congenital adrenal hyperplasia, Cushing and Addison diseases, among others ^74^. Future studies can build on this resource to explore and validate the role of endocrine signalling in human fetal brain development, study the long-term consequences of endocrine disruption and investigate the different responses to endocrine alterations across genetic backgrounds, dosing schemes and developmental stages.

**Figure 7.**
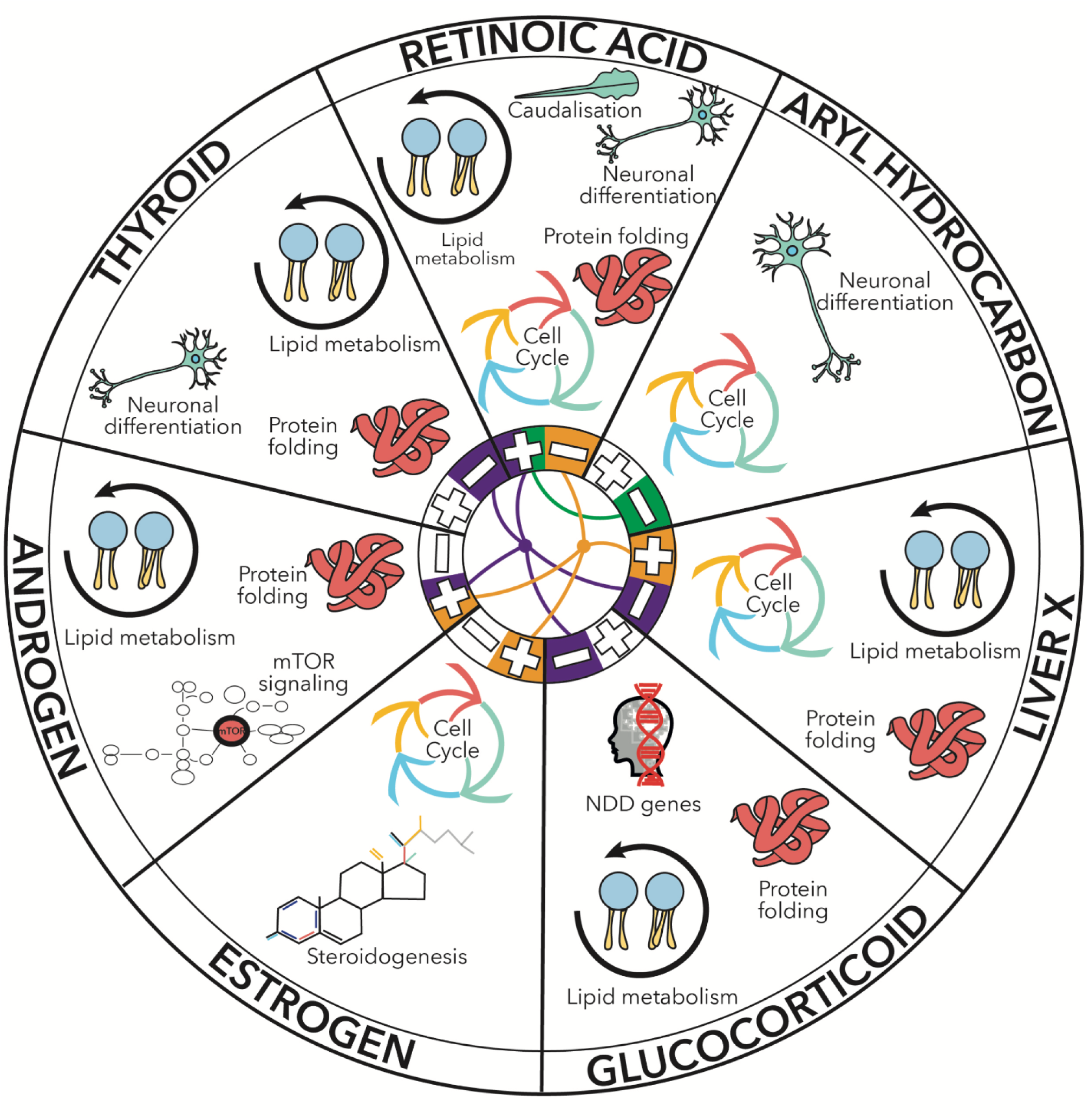
Hallmarks of endocrine signalling in human neural organoids. Schematic representation of the main molecular pathways altered by each endocrine axis in human neural organoids. The inner circle highlights the convergence across agonists (illustrated with the +) and inhibitors (illustrated with the -) of different hormonal pathways. The purple line highlights the convergence across RA and AND agonists and GC, TH, and LX inhibitors found by functional analysis of pathway-specific DEGs, DEGs overlaps and WGCNA analysis (red and blue modules); the green line highlights the convergence across RA agonist and AH inhibitor found by functional analysis of pathway-specific DEGs, DEGs overlaps and WGCNA analysis (green module). The golden brown line highlights the convergence across RA inhibitor and EST, AND, and LX agonists found by steroidomics analysis.

---

**Extended Data Figure 1.**
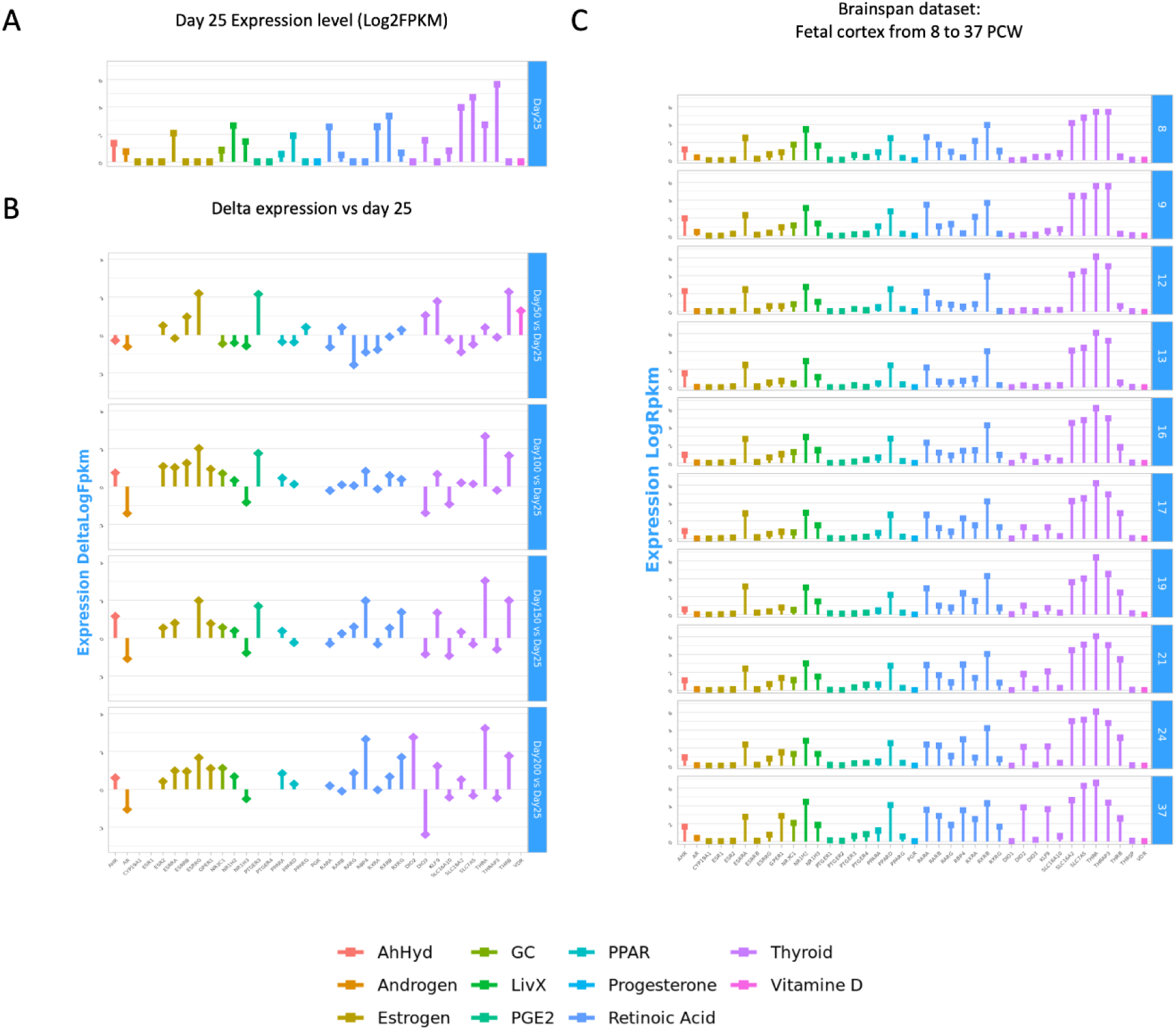
Expression levels of endocrine-related genes in neural organoids and human fetal brain (bulk transcriptomics). The analyzed pathways are: androgen, aryl hydrocarbon, estrogen, glucocorticoid, liver X, prostaglandin E2 (PGE2), peroxisome proliferator–activated receptors (PPAR), progesterone, retinoic acid, thyroid, and vitamin D. **A.** Mean log2(FPKM) expression values of selected hormonal pathway–related genes at day 25 of neural organoid differentiation from ^11^. **B.** Mean Δlog2(FPKM) expression values relative to day 25 for later stages of differentiation (days 50 to 200) in the same organoid dataset as in A. **C.** Log2(RPKM) expression values for the same gene sets in human fetal brain cortex samples from the 8th to the 37th post-conceptional week (PCW), based on data from the BrainSpan Atlas of the Developing Human Brain ^38^.

**Extended Data Figure 2.**
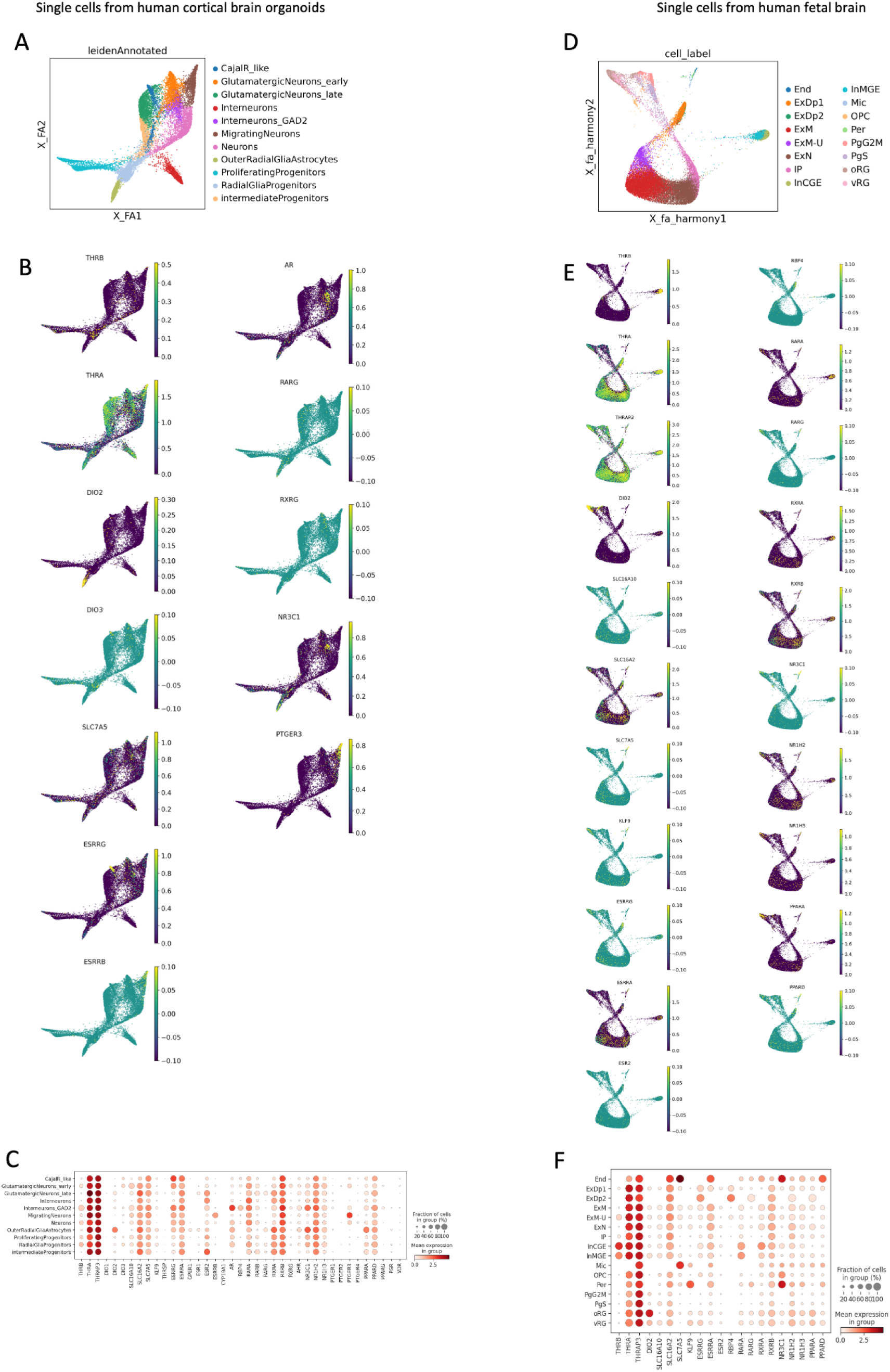
Expression levels of endocrine-related genes in neural organoids and human fetal brain (single-cell transcriptomics) for the same pathways as in Extended Data Figure 1. **A.** Force-directed graph showing cell annotations of the single-cell RNA sequencing dataset from neural organoids ^10^. **B.** Normalized expression of selected genes visualized on the force-directed graph for the neural organoid dataset. **C.** Dot plot showing mean expression levels of genes of interest across cell populations at the metacell level in the neural organoid dataset. **D.** Force-directed graph showing cell annotations of the single-cell RNA sequencing dataset from human fetal cortex ^39^. **E.** Normalized expression of selected genes visualized on the force-directed graph for the fetal cortex dataset. **F.** Dot plot showing mean expression levels of genes of interest across cell populations at the metacell level in the fetal cortex dataset.

**Extended Data Figure 3.**
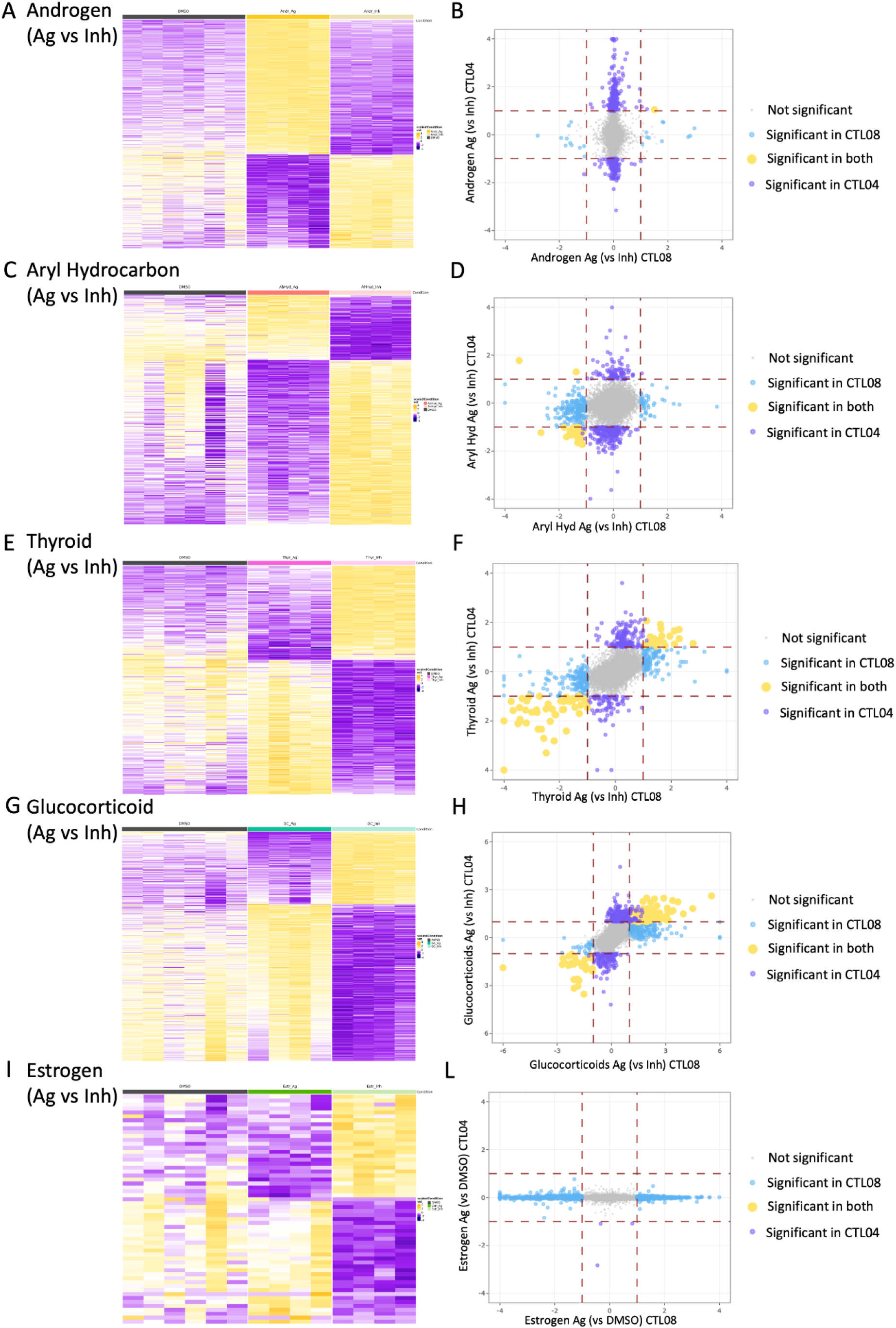
Differential expression analysis results from bulk transcriptomics in CTL04E for AND (A-B) AH (C-D), THY (E-F) GC (G-H), EST (I-L). **A,C,E,G,I.** Heatmap showing VST-normalized gene expression values (scaled by row) for differentially expressed genes (DEGs) from the Ag vs Inh comparison for each of the above-listed pathways in the female line. DMSO-treated samples are included for reference. **B, D, F, H, L.** Comparison of differential expression analysis results (Ag vs Inh) between female and male lines for each pathway. Light blue dots represent DEGs specific to the male line (CTL08), violet to the female line (CTL04), and yellow indicates shared DEGs, whether regulated in the same or opposite direction. On the axes are the log2 fold-changes. For EST (panel L) Ag vs DMSO comparison is depicted (because estrogen Inh is not available for the male line).

**Extended Data Figure 4.**
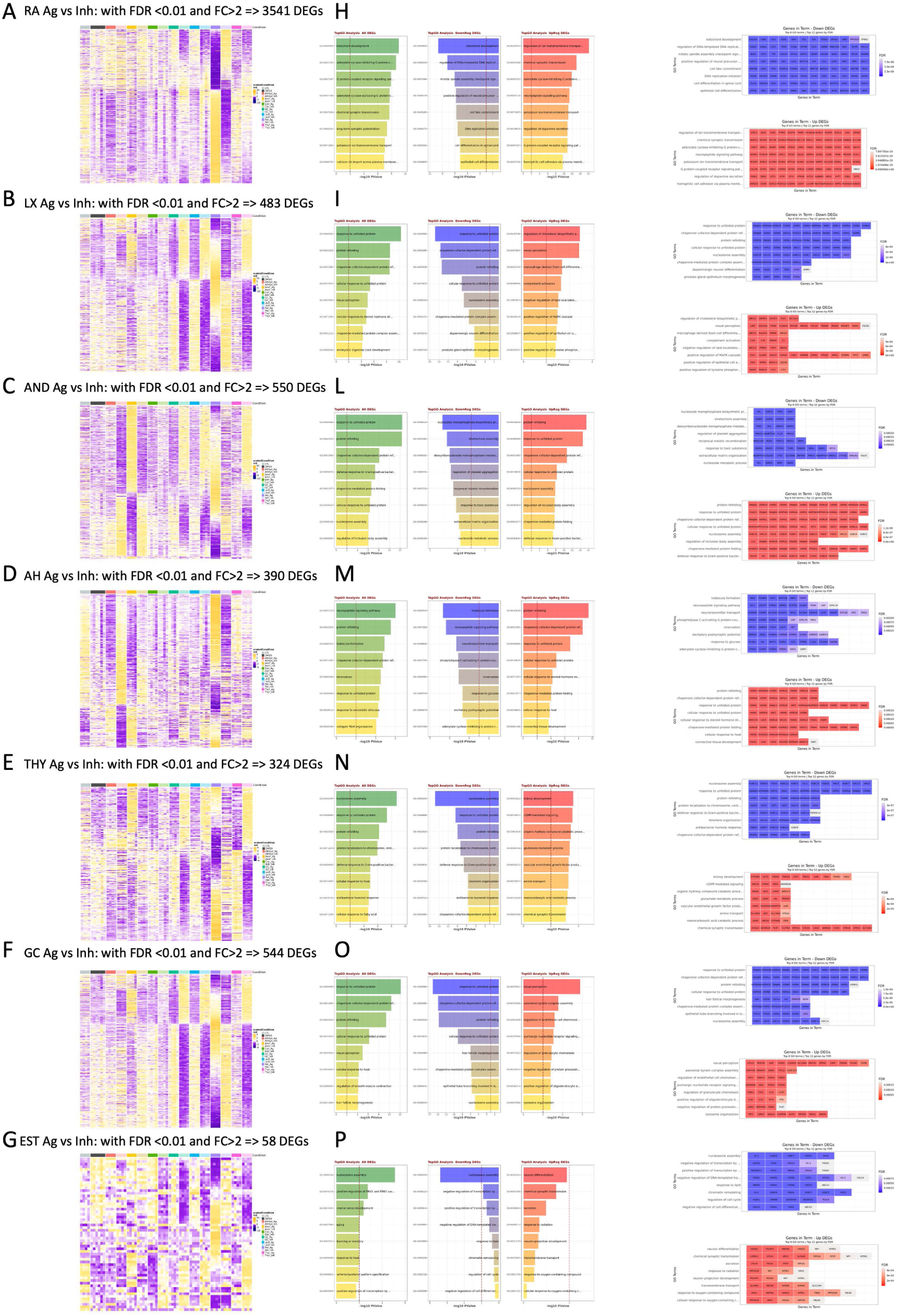
Differential expression and functional enrichment analysis results from bulk transcriptomics for all pathways in the female line. **A-G**. Heatmaps showing VST-normalized gene expression values across the complete CTL04E dataset (scaled by row) for differentially expressed genes (DEGs) from the Ag vs Inh comparison for RA (A), LX (B), AND (C), AH (D), THY (E) GC (F) and EST (G). **H-P**. Bar plots displaying the top 8 enriched Biological Process GO terms (ranked by p-value) for: all DEGs (green bars), down-regulated DEGs (blue bars), and up-regulated DEGs (red bars) for each pathway. The red vertical dashed line indicates the significance threshold at p-value = 0.01 (−log10(p-value) = 2). On the right, plots that show the top 12 DEGs (ranked by FDR) associated with each enriched term for the up- (in red) and down-regulated (in blue) gene sets.

**Extended Data Figure 5.**
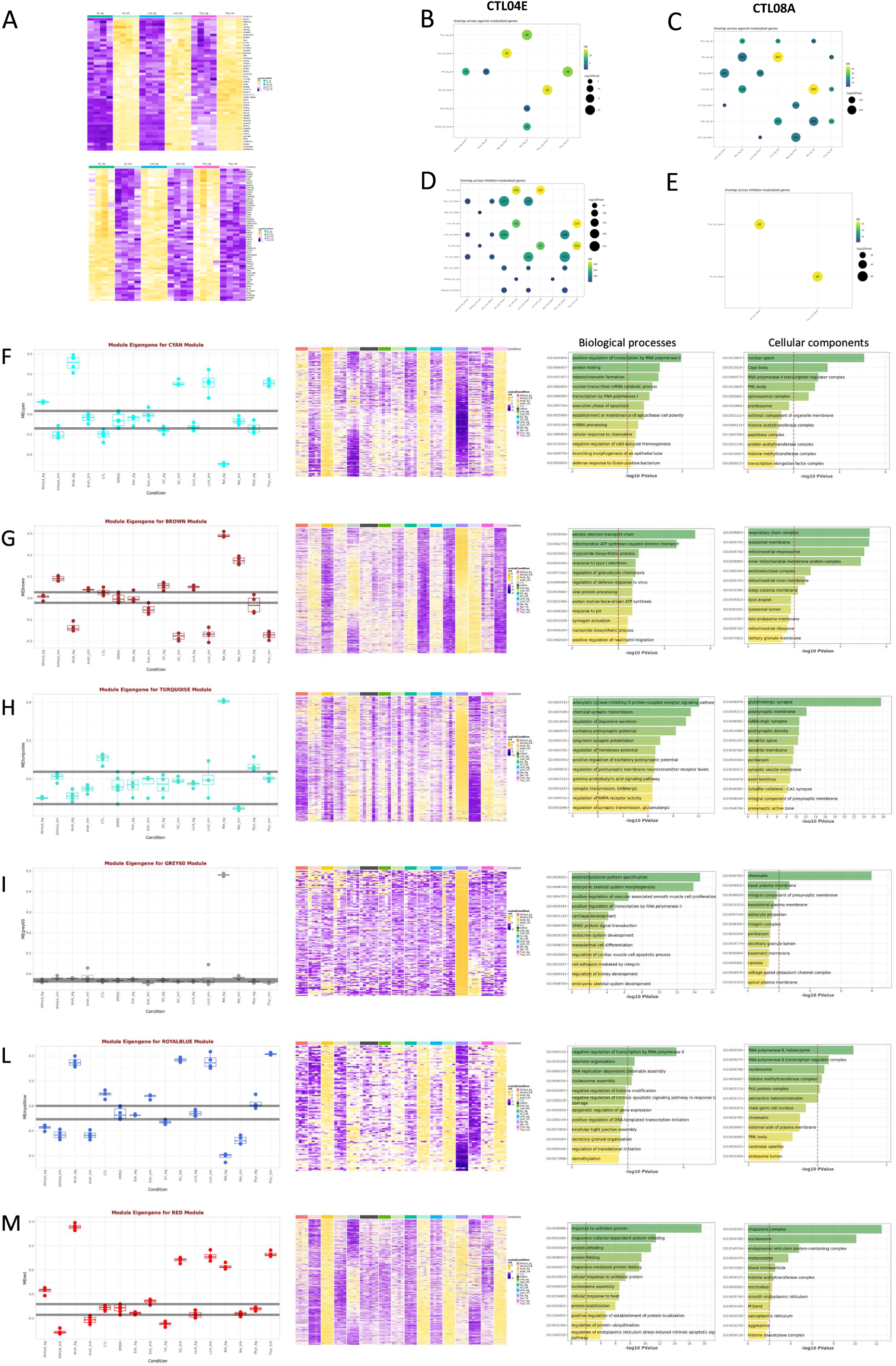
Selected examples of hormonal pathway crosstalk in neurodevelopment based on significant overlap of differentially expressed genes (DEGs) and weighted gene co-expression network analysis (WGCNA). **A.** Heatmaps of VST-normalized gene expression values (scaled by row) for the genes identified from DEGs overlap for the Ag vs Inh analysis in glucocorticoid, liverX and thyroid (upper panel DEGs upregulated by inhibitors; lower panel DEGs downregulated by inhibitors). The heatmaps show the genes across Agonists and Inhibitors of the mentioned exposures. **B-E**. Bubble plot showing DEGs overlap across the following conditions: B. Agonist exposed samples in the female cell line (CTL04), based on comparisons to DMSO; C. Agonist exposed samples in the male cell line (CTL08) compared to DMSO; D. Inhibitor exposed samples compared to DMSO in CTL04; E. Inhibitor-exposed samples compared to DMSO in CTL08A. Numbers represent shared genes; dot colour is assigned according to OR values and dot size according to p-value as shown by each legend. Significant overlaps were selected according to the following thresholds: OR > 2, p-value < 0.01, number of overlapping genes > 25. **F-M**. Analysis of the cyan (F), brown (G), turquoise (H), grey60 (I), royalblue (L), red (M) WGCNA modules. For each module: (I) left: boxplot of module eigengene values across treatments; dark grey horizontal lines represent the most extreme DMSO samples, included for reference; (II) middle, heatmap of VST-normalized gene expression values (scaled by row) for module genes across all samples; (III) right, bar plots of the top 12 significantly enriched Biological Process and Cellular Component GO terms for the module genes. Red dashed line marks the p-value threshold at 0.01 (−log10(p-value) = 2).

**Extended Data Figure 6.**
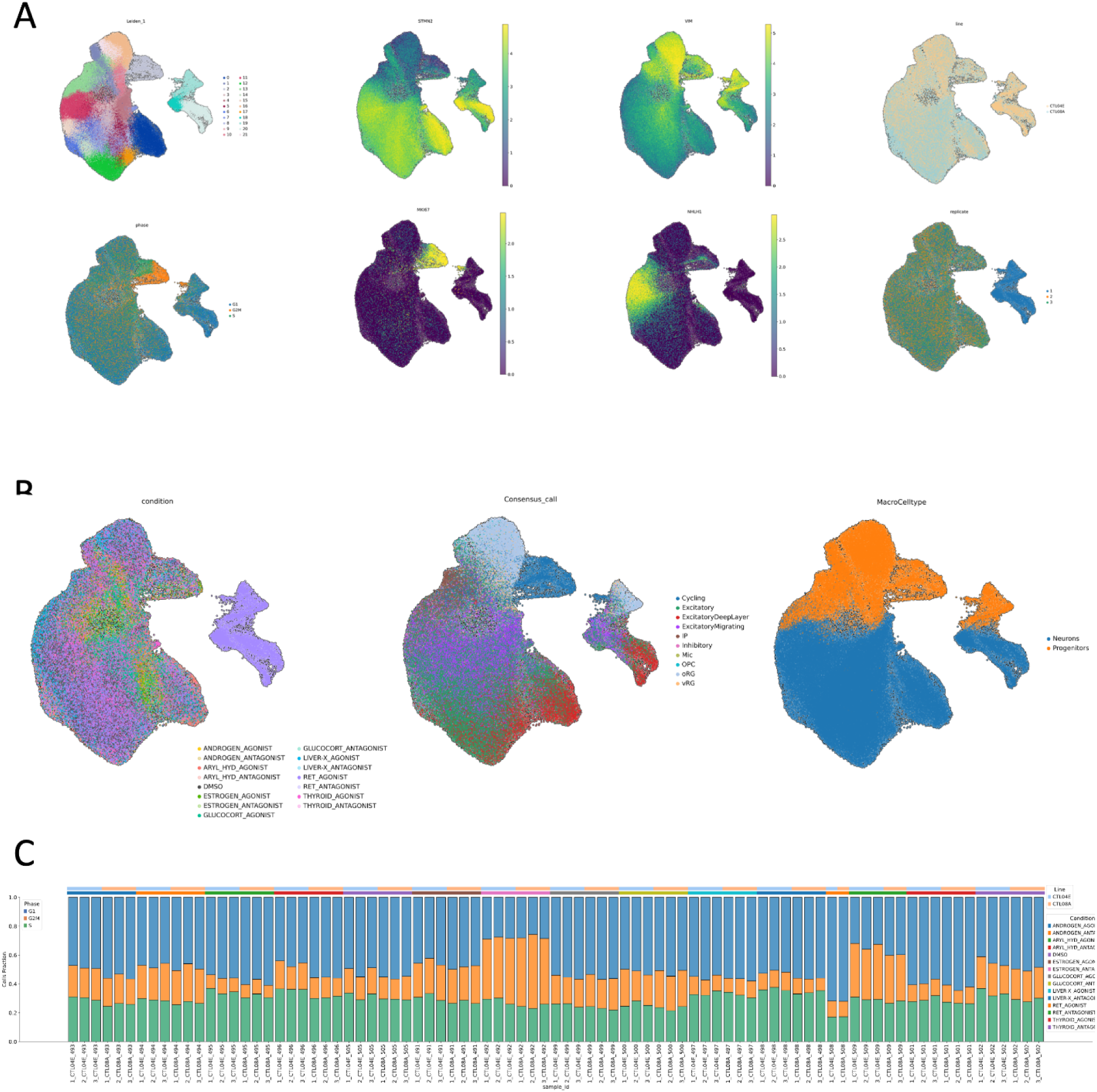
**A.** Cell embeddings in UMAPs after preprocessing, filtering. Each dot is a cell, colored by leiden cluster; cell cycle phase; expression of neurodevelopmental relevant marker (STMN2 for neuron; MKI67 for proliferating progenitors; VIM for neural progenitors; NHLH1 for intermediate progenitors); hiPSC line; experimental replicate. **B.** Cell embeddings in UMAPs after preprocessing and filtering. Each dot is a cell, colored by exposure condition; cell type label as inferred from label transfer from a single-cell RNA sequencing dataset from human fetal cortex ^39^; annotated cell types. **C.** Barplots indicating the fraction of cells assigned to the different cell cycle phases for each replicate of the different exposure conditions of neural organoids from both hiPSC lines.

**Extended Data Figure 7.**
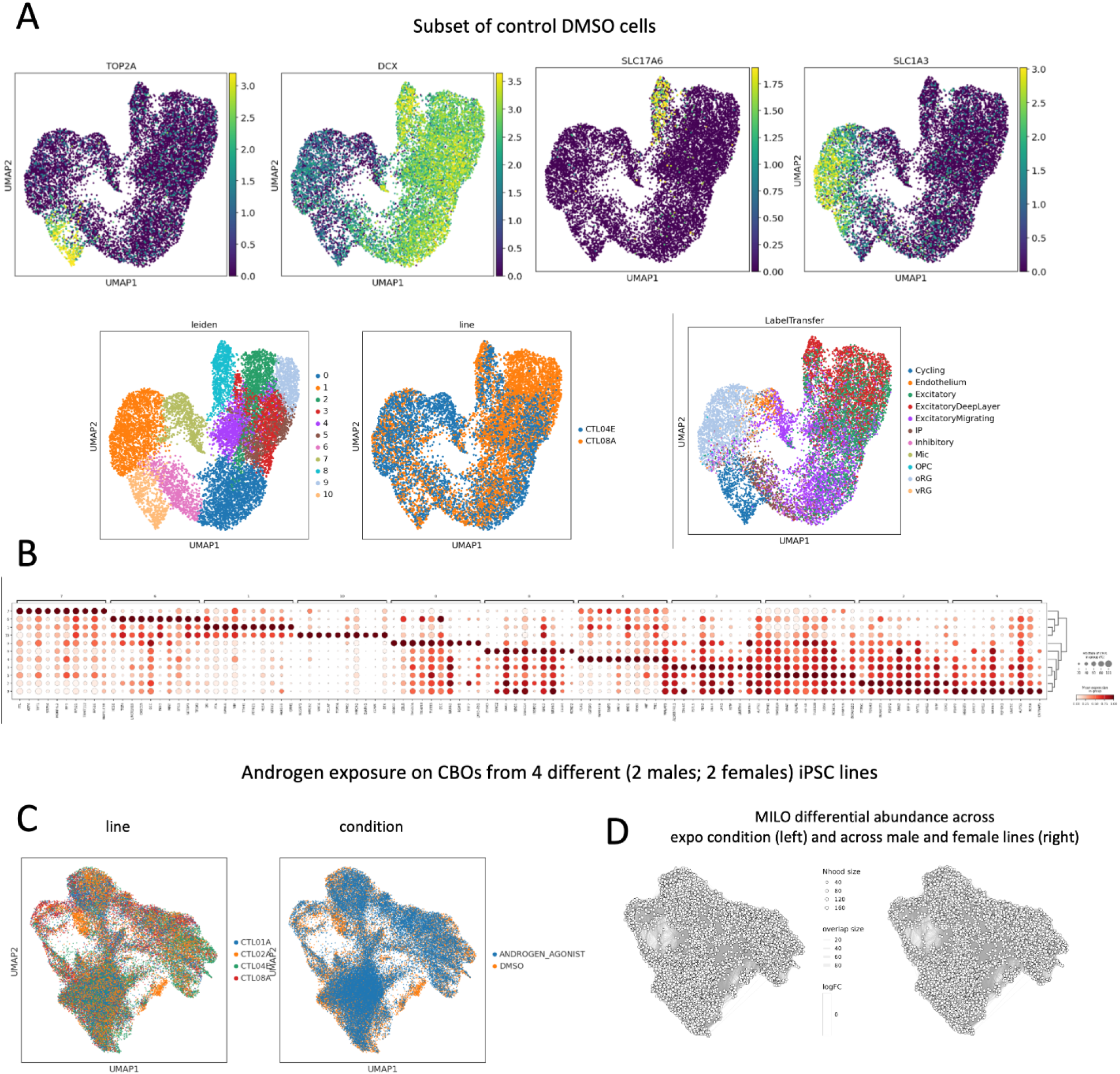
**A.** Cell embeddings in UMAPs after preprocessing, filtering and subselecting only the cells coming from negative control organoids (exposed to vehicle DMSO) of both hiPSC lines. Each dot is a cell, colored by expression of neurodevelopmental relevant marker (DCX for neuron; TOP2A for proliferating progenitors; SLC17A6 for glutamatergic neurons; SLC1A3 for radial glia) ^110^; leiden cluster; hiPSC line; cell type label as inferred from label transfer from a single-cell RNA sequencing dataset from human fetal cortex ^39^. **B.** Dot plot of gene expression for some of the relevant markers used in the annotation. Size of dots is proportional to the number of cells expressing a marker, and color encodes the mean expression in the group (scaled log-normal counts). **C.** Cell embeddings in UMAPs after preprocessing, filtering and subselecting only the cells coming from the independent experiment where neural organoids from four hiPSC lines (two males and two females) were exposed to AND agonist. Each dot is a cell, colored by hiPSC line; exposure condition. **D.** Differential abundance graph ^53^ across exposure condition (AND vs DMSO) and across male and female hiPSC lines. The basis for the visualization is the UMAP. Dots represent groups of similar cells; color code indicates enrichment for the indicated categories (no significantly enriched neighborhoods were identified, spatial FDR < 0.1)

**Extended Data Figure 8.**
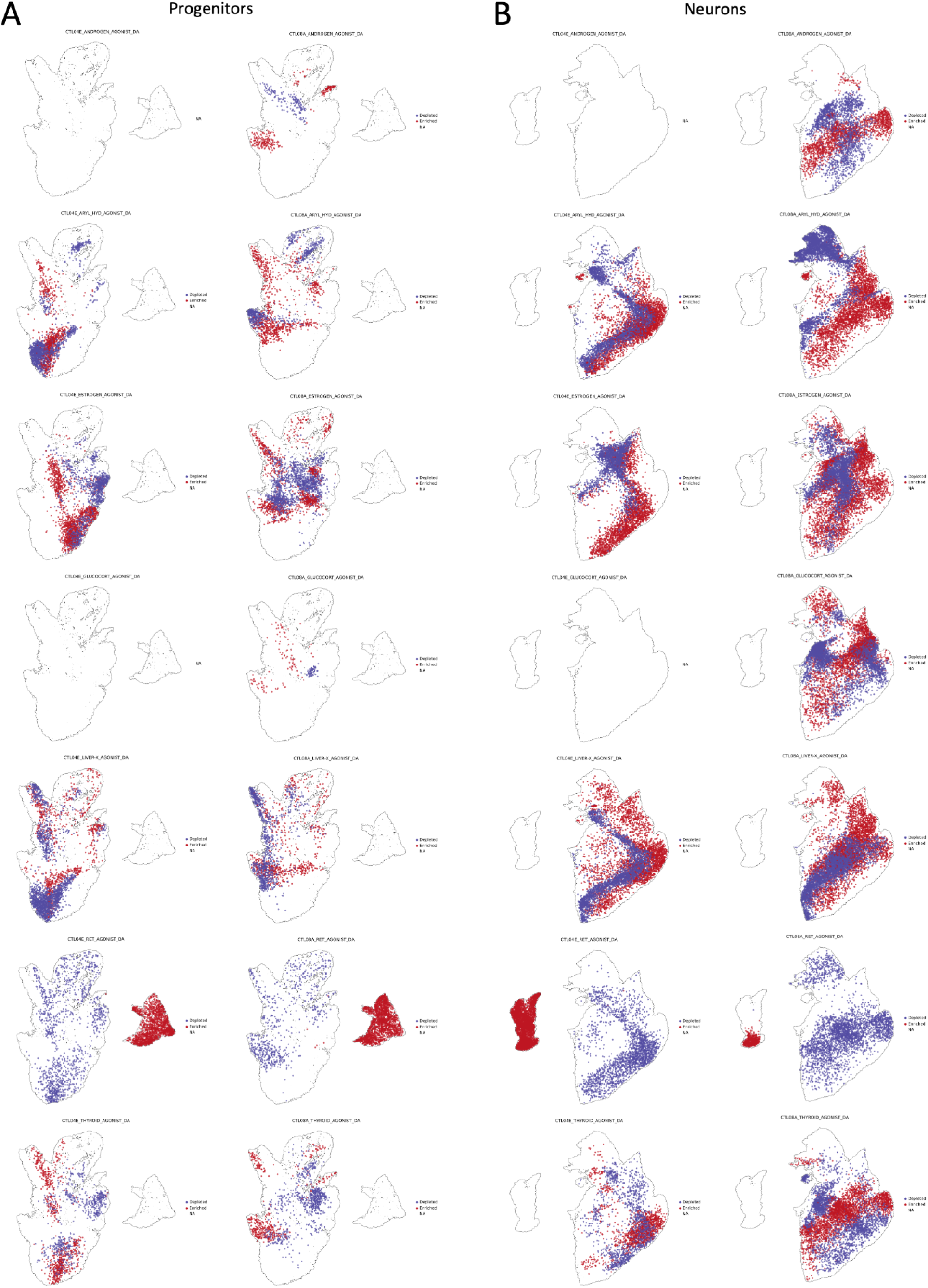
Differential abundance graph ^53^ between agonist and inhibitor of each endocrine pathway. The analysis was performed independently on neural organoids from the male and female hiPSC lines, and on progenitors (**A**), and neurons (**B**). The basis for the visualization is the UMAP. Dots represent cells; color code indicates enrichment/depletion for the indicated categories (only shown for significantly enriched neighborhoods, spatial FDR < 0.1)

**Extended Data Figure 9.**
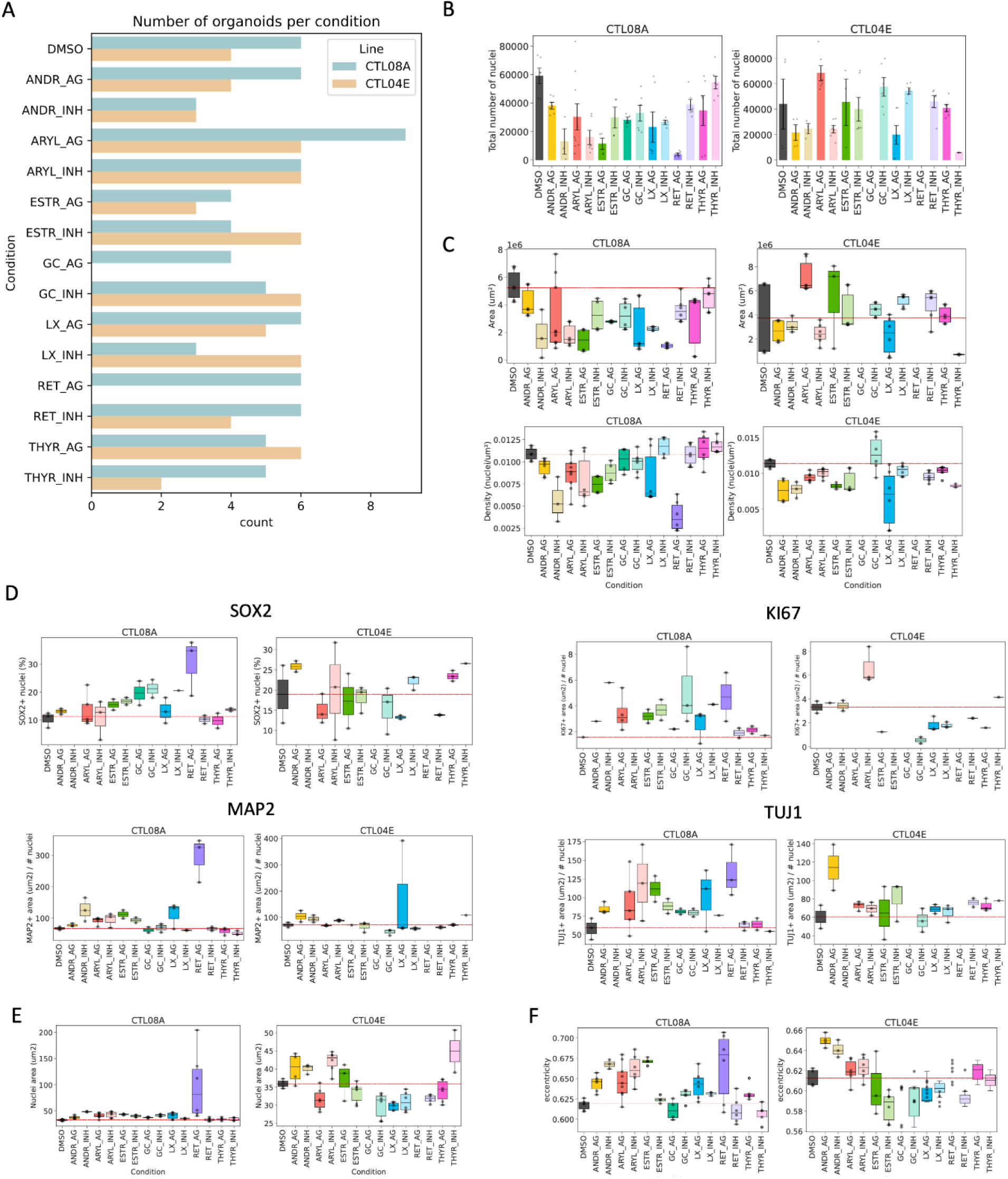
Organoids morphological descriptors and expression of cell-type specific markers across hormonal conditions. **A.** Barplot showing the number of organoids used for the analysis of immunofluorescent markers after manual quality control of tissue integrity. **B.** Total number of nuclei segmented per organoid and used for the analysis across conditions and lines. Error bar refers to the standard error of the mean. **C.** Area (top) and density of nuclei (bottom) per organoid. **D.** Top: quantification of the percentage of nuclei positive to SOX2 and KI67 per organoid across conditions. Bottom: ratio between area positive to MAP2 and TUJ1 and number of total nuclei. **E.** Mean nuclei area per organoid. F. Mean nuclei eccentricity per organoid. For all the boxplots shown, the solid black line of each box is the median of the distribution; whiskers show the first and third quartiles while the red dotted line is displaying the median of the DMSO. Only organoids that passed the quality check for tissue integrity and staining were kept for the plot.

**Extended Data Figure 10.**
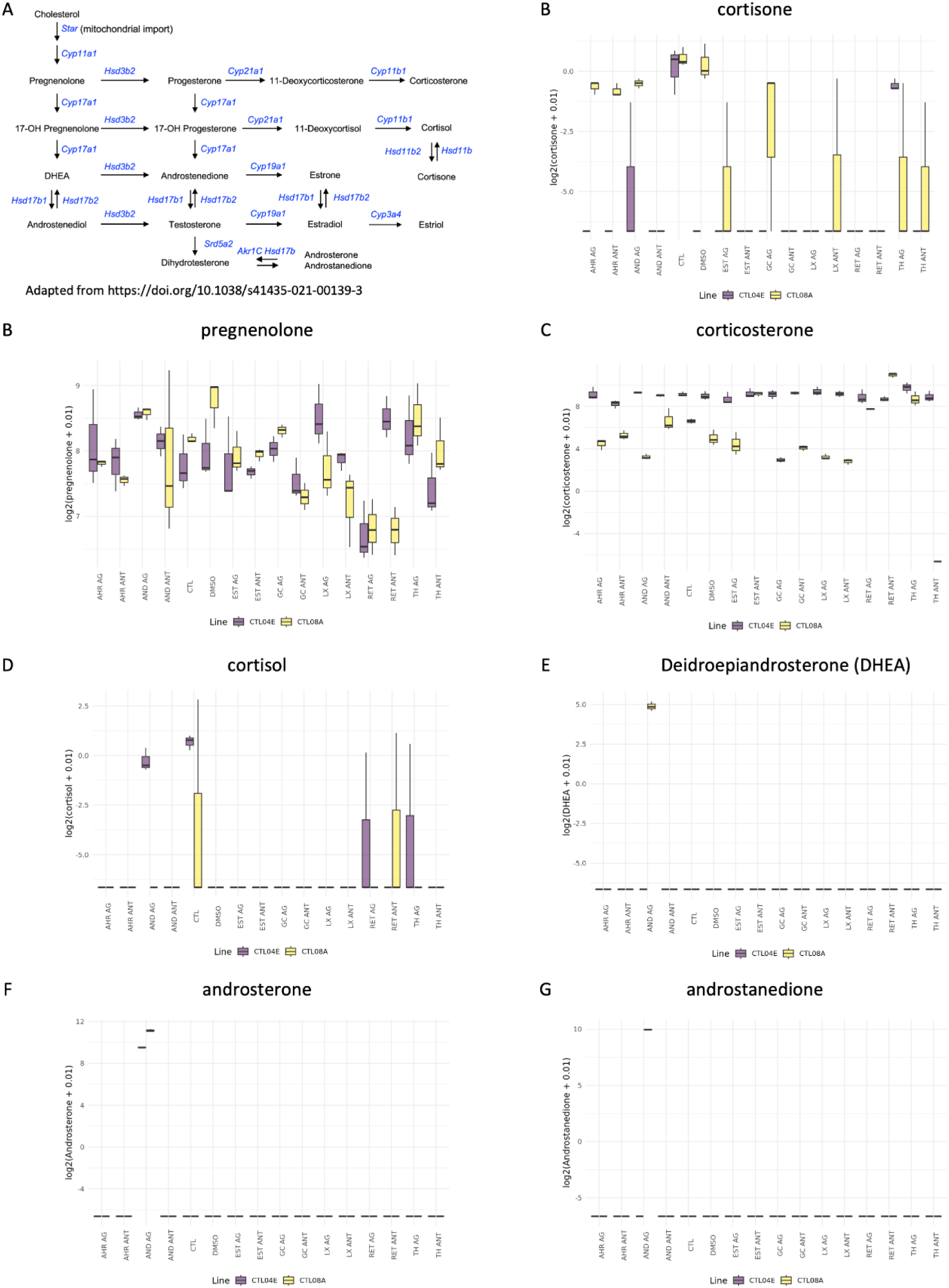
Steroidogenesis profiling in neural organoids upon endocrine perturbations **A.** Schematic representation of the steroidogenesis pathway including the steroid metabolites quantified in neural organoids and the enzymes catalising the metabolic reactions**. B-G**. Boxplots of the log2 values of the measured steroid metabolites in neural organoids from both hiPSC lines exposed to endocrine agonists and inhibitors.

## Methods

### Human iPSCs maintenance

Human iPSC lines were cultured under feeder-free conditions at 37 °C, 5 % CO2 and 3 % O2 on matrigel-coated plates in TeSR/E8 medium (Stemcell Technologies, 05990) supplemented with 100 U/mL penicillin and 100 ug/mL streptomycin (ThermoFisher, 15140122). Media changes were performed daily and hiPSCs were routinely passed 1:10 with ReLeSR (Stemcell technologies, 05872) upon reaching 70% confluency. When single cell dissociation was needed, Accutase (Sigma-Aldrich, A6964) was used instead of ReLeSR, and 5uM ROCK inhibitor Y-27632 (Tocris, 1254) was added to the medium during the first 12 hours to enhance cell survival. To comply with the guidelines for best practices for human stem cell (ISSCR guidelines: [link](https://static1.squarespace.com/static/611faaa8fee682525ee16489/t/62ed69b184e2ed258e6eb7e4/1659726257773/isscr-guidelines-for-stem-cell-research-and-clinical-translation-2021.pdf)) and *in vitro* neurotoxicology ^75^, hiPSCs banks have been established and validated in compliance with the requirements of hPSCreg ^76^. Two hiPSC lines (1 male and 1 female) were used, namely WTSIi018-B (here named CTL08, alternative name KOLF_2) one of the best characterized iPSC lines, recently proposed as a reference for standardized disease modeling projects ^77^, and UMILi024-A (here named CTL04), both previously used by us in ^10,11^, and already registered in the hPSCreg registry ^76^. HiPSCs were routinely tested for Mycoplasma by PCR assay and used within 3-7 passages from thawing of the cell bank vial. All hiPSC lines employed were approved by the ethical committee of the University of Milan.

### Preparation of test chemicals and mixtures stock solutions

All the chemicals used in this work were dissolved in DMSO to obtain a 10 mM (1 x 10^-2^ M) stock solution unless differently specified in the material datasheet and stored at –80 °C in ambered glass vials. Before the beginning of chronic exposure, stock solutions were decimally diluted to obtain working solutions 1 x 10^3^ the final concentration of use to avoid exceeding 0.1% DMSO concentration in medium. Working solutions were aliquoted upon preparation into single use tubes to avoid multiple freeze/thaw cycles. The concentration of use of hormonal receptors agonist and inhibitors was agreed by the members of the ENDpoiNTs consortium ([link](https://endpoints.eu/)). The list of the test chemicals employed in the project according to previous evidence from human biological models can be found in Table 1. Both stock and working solutions were kept at -80 °C for a maximum of 1 year since preparation.

### Neural organoids differentiation

Neural organoids were generated according to the protocol originally published by ^40^, with minor modification to improve efficiency as previously published by us in ^6,10,78^. Briefly, hiPSCs were dissociated with Accutase (Sigma, A6964) as previously described and resuspended in TeSR/E8 supplemented with 5uM Y-27632 (Tocris, 1254) to reach a final concentration of 2 x 10^5 cells/ml. 100uL/well of cell suspension were seeded into PrimeSurface, U-bottom 96 well plates (SystemBio, MS9096UZ), centrifuged for 3 minutes at 150 RCF to promote aggregation of embryoid bodies and incubated for 60 h at 37 °C, 5% CO2, 3% O2. Next, first media change was performed (DIV 0), replacing TeSR/E8 with neural induction medium 1 containing 80% DMEM/F12 medium (Gibco, 11330057), 20% Knockout serum (Gibco, 10828028), Non-essential amino acids 1:100 (Sigma, M7145), 0.1 mM cell culture grade 2-mercaptoethanol (Gibco, 31350010), GlutaMax 1:100 (Gibco, 35050061), penicillin 100 U/mL and streptomycin 100 μg/mL (ThermoFisher, 15140122), 5 µM Dorsomorphin (Sigma, P5499) and 10 µM TGFβ inhibitor SB431542 (MedChem express, HY-10431). From this moment onward, cultures were grown in normal oxygen conditions (21% O2). Media changes were performed daily for the subsequent 4 days and, on DIV 5, neural induction medium was substituted with with complete neurobasal medium, composed of neurobasal medium (Gibco, 12348017) (in particular we used neurobasal without phenol red to avoid its estrogenic activity as a confounder), B-27 supplement without vitamin A (1:50, Gibco, 12587001), GlutaMAX (1:100, Gibco, 35050061), penicillin at 100 U ml−1 and streptomycin at 100 μg ml−1 (Thermo Fisher, 15140122) and 0.1 mM cell culture-grade 2-mercaptoethanol solution (Gibco, 31350010) supplemented with 20 ng ml−1 FGF2 (PeproTech, 100-18B) and 20 ng ml−1 EGF (PeproTech, AF-100-15). On day 12, organoids were transferred by pipetting with cut-end pipette tips from 96-well to 9-cm ultra-low-attachment dishes (System Biosciences, MS-90900Z) and placed on a standard orbital shaker (VWR Standard Orbital Shaker, Model 1000). From day 12 onward, medium changes were performed every other day. On day 23, FGF and EGF were replaced with 20 ng ml−1 BDNF (PeproTech, 450- 02) and 20 ng ml−1 neurotrophin 3 (PeproTech, 450-03) to promote differentiation of neural progenitors. From day 42 onward, complete neurobasal medium without BDNF and NT3 was used, performing medium changes every other day. Following the indications from ^8^, for each assays performed, we included at least 3 replicates per exposure condition. If not specified differently, replicates are different organoids from the same differentiation round. For the bulk transcriptomics profiling of CTL08 we had to perform two different rounds of differentiations, which are indicated as metadata (SeqRun 20210310 and 20210724 for the first differentiation round, SeqRun 20220422 for the second differentiation round) in all the relevant plots included in the GitHub repository, and used as covariate to perform differential expression analysis.

### Exposure to test compounds

Nomenclature specifics: while we acknowledge that in the field of toxicology and pharmacology, an agonist is a chemical that binds to a cellular receptor and activates it to produce a biological response; an antagonist is a chemical that binds to a receptor but does not activate it; an inhibitor is often used more broadly and can refer to a substance that decreases the rate of a specific biochemical reaction, typically by binding to an enzyme (Goodman & Gilman’s: The Pharmacological Basis of Therapeutics, 14th Edition), in this study we use these terms in a functional sense, to indicate the overall effect on the specific endocrine pathways. Therefore, for the purposes of this work, "agonist" simply refers to a compound that causes the activation of an endocrine pathway, while "antagonist" or "inhibitor" refers to a compound that causes the inhibition of that pathway.

During the first 12 days of neural organoids differentiation, exposure was performed by dissolving the necessary amount of single use working solution to a fresh aliquot of media right before media change. This was done to avoid chemical degradation and ensure proper chemical concentration was equally administered at every media change. From day 12 onwards, to minimize the issues related to precipitation of poorly water-soluble chemicals in the medium during its storage into intermediate vessels, chemicals were added directly in the culture dish as follows. Briefly, spent medium was removed while tilting the plate to collect the organoids on one side. Next, to avoid contact between organoids and concentrated vehicle or test chemical, the desired amount of working solution was added to the opposite side of the plate with a Gilson pipette and immediately flushed with fresh medium to disperse homogeneously the treatment.

### Immunofluorescence

Organoids were collected on DIV 50 (+/- 3 days), washed three times in PBS and fixed overnight at 4 °C in 4% paraformaldehyde/PBS solution (SantaCruz, sc-261692). After rinsing with PBS twice, samples were embedded in 2% w/v low melting agarose/PBS (3-4 organoids/block) and dehydrated overnight in 70% v/v ethanol/water solution. Dehydration and paraffin embedding were performed using Sakura VIP6AI.

To increase the throughput of the assay while reducing the technical variability between samples, single paraffin-embedded blocks were assembled into paraffin-encased Tissue Microarrays (TMAs) containing up to 6 experimental conditions. Briefly, paraffinized blocks were heated to 65 °C to melt the excess paraffin and laid in a 3x2 grid into disposable plastic molds. A small fragment of China ink-dyed agarose block was also placed at the top right corner of the mold for spatial reference. The chamber was then filled with liquid paraffin and left to solidify on a cooled surface. 3 µm sections were cut at the microtome and stored in a dark and dry environment for maximum 2 months before processing.

Deparaffinization and rehydration was achieved by consecutive passages of 5 minutes each in the following solutions: 2X xylene, 2X 100% ethanol, 2X 95% ethanol, 80% ethanol and 2X ddH2O. Antigen retrieval and tissue permeabilization was performed by incubating the sections for 45 minutes at 95 °C with 10mM Sodium citrate buffer Ph.6 (WVR chemicals, 27833.294P) + 0,05% Tween 20 (Sigma, P1379), followed by equilibration at RT for at least 2 hours. Slides were then rinsed in TBS and incubated for 60 minutes with blocking solution containing 5% normal donkey serum (Jackson ImmunoResearch, 017-000-121) and 0.3% Triton X-100 (Sigma, T8787) in TBS. Primary antibodies were diluted in blocking solution and incubated overnight at 4 °C. On the following day, secondary antibodies conjugated with Alexa Fluor 488, 555 or 647 diluted 1:400 in PBS + 1 μg ml^−1^ DAPI solution were applied to the sections for 1 hour. After each incubation 3 x 5 minutes washing steps with TBS buffer were performed. After a final rinse in deionized water, slides were dried and mounted using Mowiol mounting solution. All the images were acquired at 20X magnification using ZEISS Axio Scan.Z1 in a single batch of acquisition. The list of used antibodies and their concentration is reported in Table 2.

### RNA extraction and bulk sequencing

Neural organoids were harvested at DIV 50 into 2ml Eppendorf tubes and washed three times with 1.5 ml PBS to remove medium residues before being snap-frozen and stored at -80 °C for a maximum of 1 month before RNA extraction. For each of the replicates, 1 organoid was collected unless size constraints required the pooling of multiple organoids (i.e. Retinoic Agonist condition group, 3 organoids for each replicate). After thawing the pellets on ice, RNA was extracted using the PureLink RNA Mini kit (Thermofisher, 12183025) and eluted in 30ul of RNAse free water. On-column DNAse I treatment was performed with RNAse-free DNase set (Qiagen, 79265) according to protocol recommendations. Purified RNA was quantified with Nanodrop One (Thermofisher) to estimate the concentration range, while the RNA Integrity Index (RIN) was measured by TapeStation analysis. A RIN >7.5 (scale 1-10) was chosen as a cutoff for further sample processing. Samples with RIN <7.5 were discarded and RNA was re-extracted from backup pellets.

RNA concentration was then normalized, and cDNA libraries were prepared using Illumina Stranded mRNA Prep with polyA-capture technology with an initial RNA input of 100 ng/sample. Samples were sequenced in paired-end configuration with a target coverage of 35 million reads/sample.

### Quantification of steroid species in neural organoids

Steroid hormones were extracted using a modified method from ^79^, excluding the deconjugation step and optimizing the procedure for neural organoids analysis. Since the method from ^79^ focused on liver and plasma, performance was re-evaluated to confirm its suitability for extraction and analysis in the new matrix. Specifically, 100 μL of ice-cold water was added to the tubes, and the samples were homogenized using a Precellys mill (Bertin Instruments, Montigny-le-Bretonneux, FR) for one cycle of 10 sec at 6500 rpm. The samples were placed in an ice-cold ultrasonic bath for 5 min. Subsequently, 12 μL of each homogenate was transferred to a 96-well plate for the Bradford protein assay. The remaining homogenates were added with 50 μL steroid hormone internal standard (IS). IS solutions were prepared to maintain organic solvent content below 1%. Samples were homogenized again with the Precellys homogenizer (Bertin Instruments, Montigny-le-Bretonneux, FR) for 10 s at 6500 rpm, followed by the addition of 200 μL of acetonitrile (ACN) and sonication for 15 min. Then, 1400 μL of 2% formic acid (FA) water (v/v) was added, and the mixture was centrifuged at 13000 rpm for 10 min. The supernatant was then processed through solid phase extraction (SPE) and transferred to a 96-well SPE plate (VersaPlate, Agilent Technologies, Santa Clara, CA, U.S.) with Bond Elut Plexa cartridges (30 mg), which were conditioned with 1 mL of methanol (MeOH) and 1 mL of 0.5% FA water (v/v). After loading the supernatant, the cartridges were washed and dried, and the samples were eluted with 1 mL of 100% MeOH. The eluates were evaporated in a CentriVap vacuum concentrator (Labconco, Kansas City, MO, U.S.) at 40 °C, then reconstituted in a 75 μL MeOH/water (1:1) mix and shaken for 20 min. The plate was sealed and centrifuged for 2 min (13000 rpm) for further LC-MS/MS analysis using an ExionLC system coupled with a SCIEX Triple Quad 6500+ mass spectrometer (SCIEX, Framingham, MA, U.S.) equipped with electrospray ionization (ESI) source operating in positive mode for steroid hormones and dansylated estrogens. The extracts were analyzed twice: first for steroid hormones and then for steroid sulfates. Source parameters, MRM settings, and retention times for steroid hormones including estrogens are reported in Table 3. Samples were randomized before analysis, and the calibration curve standards were injected after every 10 samples and two times during each run. Additionally, three procedural blanks (extracted alongside the samples) and three solvent blanks were injected at the beginning and end of the sequence to monitor potential interference and evaluate the extraction efficiency. LC-MS/MS raw data acquisition and processing for steroid hormones were conducted using AB SciexOS Analyst version 1.6.2 (SCIEX, Framingham, MA, U.S.). The compound concentrations were estimated based on internal standards and external calibrators.

### Bradford assay for protein concentration analysis

Protein concentration in organoids homogenates was determined using the Bradford protein assay. Prior to measurement, 12 µL of each CBO homogenate was diluted 1:20 in water to ensure readings fell within the linear range of the assay. Then, 10 µL of each diluted sample was mixed with Bradford reagent (Bio-Rad), and absorbance was measured at 595 nm after incubation for 5–10 minutes at room temperature. A standard curve was prepared using serial dilutions of bovine serum albumin (BSA; 0–1 mg/mL), and protein concentrations were calculated based on the standard curve. All measurements were performed in duplicate. Protein content (average between the two measurements) was used to normalize sample concentration input for downstream analyses. Table with normalized measurement for molecules detected in at least one condition are available at [[GitHub link](https://github.com/GiuseppeTestaLab/noha/blob/main/4_steroidomics/Normalized_Counts_Neuralorganoids_Steroidomics.xlsx)].

### Comparison of steroid concentrations with WGCNA modules

To perform an analysis across scales, we examined the relationship between the WGCNA modules from the bulk transcriptomics and the levels of compounds measured by steroidomics. Only steroids that were quantified in at least 8 samples were kept. Four steroids (pregnenolone, testosterone, androstenedione, estrone (E1)) had Spearman correlation coefficient > 0.6 and p-value < 0.01 with at least one WGCNA module, and were thus visualised in scatterplots [[GitHub link](https://giuseppetestalab.github.io/noha/Steroidomics_WGCNA.html)].

### Hormonal-related gene exploration in bulk and scRNAseq datasets

Literature-curated gene signatures related to various hormonal pathways were explored across four transcriptomic datasets. The pathways analyzed include androgen, aryl hydrocarbon, estrogen, glucocorticoid, liver X, prostaglandin E2 (PGE2), peroxisome proliferator–activated receptors (PPAR), progesterone, retinoic acid, thyroid, and vitamin D. The two bulk RNA-seq datasets used were the BrainSpan Atlas of the Developing Human Brain, downloaded from [link](http://www.brainspan.org/static/download.html), and the organoid dataset from ^11^. The two scRNA-seq datasets were the fetal brain cortex dataset from ^39^ and the organoid dataset from ^10^.

For bulk RNA-seq data, expression values (log2FPKM or log2RPKM) of the selected gene signatures were retrieved, the mean expression was calculated for each developmental stage within each dataset, and results were visualized using lollipop plots. For scRNA-seq datasets, raw counts and cell annotations were retrieved, and normalized gene expression patterns were visualized in force-directed graph plots. Furthermore, to enable a more robust exploration of gene expression across cell populations, we generated metacells using SEACells, following the official tutorial with default parameters, except for setting the metacell resolution to one metacell per 60 single cells (instead of the canonical 75) for the organoid dataset. Gene expression within metacells was visualized using dot plots.

### Bulk RNA data analysis

Unless otherwise specified, all subsequent analyses were performed on exposed organoids derived from both cell lines (CTL04 and CTL08). If different criteria were applied to the two lines, this is explicitly stated. All code used for the analyses is available at [[GitHub folder link](https://github.com/GiuseppeTestaLab/noha/tree/main/1_bulkRNASeq)]. The bulk RNA-seq analyses were executed using the Docker container “testalab/downstream:EndPoints-1.1.5”, and all R package versions were managed using renv v.0.15.4. R v.4.2.1 was used for all bulk RNASeq analyses, unless differently specified.

### Read alignment

FASTQ data were quantified at the gene level using Salmon v1.3.0 ^80^. Human Gencode Release 35 (GRCh38.p13) was used as a reference for transcript count quantification and gene annotation, retrieved with biomaRt v2.54 ^81^.

### Quality control

Several quality control metrics were calculated and evaluated for each line on the aggregated count matrix of all samples, along with exploratory plots to assess dataset quality thoroughly.

Following appropriate filtering to remove lowly expressed genes and retain only protein-coding genes, principal component analysis (PCA) was performed using the prcomp() function in R. The first two principal components were visualized and inspected for the presence of outlier samples and potential batch effects. Based on these analyses, some low-quality CTL08 samples were excluded from downstream analysis. In particular, all three replicates exposed to the estrogen inhibitor were removed (thus this condition is absent for this line in the results), along with two DMSO-treated control replicates, one replicate exposed to the retinoic acid inhibitor, and one to the glucocorticoid inhibitor. Furthermore, a batch effect was observed in the CTL08 line, where each organoid differentiation batch had been sequenced in a different sequencing run. As a result, the sequencing run variable was used as a covariate to correct for this batch effect in differential expression analysis. After quality control and filtering, 51 CTL08 and 66 CTL04 samples were retained for downstream analyses.

### Literature-curated gene signature exploration

For both cell lines, we performed a supervised exploration of gene sets manually curated from the literature, focusing on genes relevant to the biological context of the samples under study. These included markers of distinct neural lineage populations, hormone-regulated target genes, metabolic genes, and genes related to hormonal receptor activity. Normalization and calculation of FPKM and log2 FPKM values were carried out using edgeR v3.40.2 ^82^. Log2 fold-change (logFC), was computed using SEtools v1.12.0 ^83^, with DMSO-treated organoids used as the control condition. Heatmaps were generated using Sechm v1.6.0 ^84^ to visualize gene expression patterns, displaying either row-scaled log2 FPKM or log2 FC values for the selected gene sets across samples.

### Differential gene expression analysis (DEA)

Differential expression analyses were performed by comparing each agonist to its corresponding antagonist (inhibitor) for each pathway, as well as by comparing each treatment condition to DMSO-treated organoids.After selecting protein-coding genes, expression thresholds were applied to exclude non-expressed or lowly expressed ones: genes were retained if they had at least 30 reads in ≥6 samples for the CTL04 line and ≥10 samples for the CTL08 line. This filtering resulted in 14,305 and 14,850 genes retained for downstream testing in the two lines, respectively.

Differential expression analysis was then conducted using DESeq2 v1.38.3 ^85^. This included estimating size factors and dispersions, followed by variance stabilizing transformation (VST) of the count data. In the case of CTL08 samples, the sequencing run variable was included as a covariate in the statistical model to correct for batch effects. LogFC shrinkage was applied using the ashr method ^86^ to reduce noise from low-count genes. Genes were considered differentially expressed (DEGs) if they met the thresholds of FDR < 0.01 and |log2(FC)| > 1. False discovery rates (FDRs) were calculated using the Benjamini–Hochberg multiple testing correction method. The number of DEGs, separated into upregulated and downregulated genes, was quantified for each comparison, and their VST-transformed expression values were visualized using heatmaps as described above.

### Functional enrichment analysis

Functional enrichment analysis of Gene Ontology (GO) categories, Biological Process (BP), Molecular Function (MF), and Cellular Component (CC) was performed for each comparison using topGO v2.50.0 ^87^. Genes were separated into up- and downregulated sets, and enrichment was calculated using the ‘weight01’ algorithm, Fisher’s exact test, and a node size of 15. A p-value < 0.01 and enrichment > 2 defined significantly enriched GO terms.

Additionally, complementary functional enrichment analyses are available in the repository [[GitHub folder link](https://github.com/GiuseppeTestaLab/noha/tree/main/1_bulkRNASeq/4_FunctionalAnalysis)], performed using EnrichmentBrowser v2.28.2 ^88^, which includes a broader selection of gene sets and alternative statistical approaches. For network-based enrichment, pathway information was retrieved from the KEGG ^89^ and Reactome ^90,91^ databases. For gene set-based enrichment, gene sets were sourced from the Molecular Signatures Database (MSigDB) ^92,93^, specifically the H (hallmark) collection, and from the Enrichr database ^94^, using the “DisGeNET” ^95^, and “HumanCyc_2016” ^96^ libraries.

For network-based analyses, the selected statistical methods were GGEA (Gene Graph Enrichment Analysis) ^97^ and DEGraph (Differential Expression Testing for Gene Graphs) ^98^. For gene set-based enrichment, CAMERA (Correlation Adjusted MEan RAnk) ^99^ and PADOG (Pathway Analysis with Down-weighting of Overlapping Genes) ^100^ were employed. All functions were executed using default parameters.

### Analysis of DEG overlap between and within cell lines

Common DEGs between the two cell lines were visualized using scatter plots (ggplot2), displaying log2 fold changes and significance for each pathway individually to highlight genes that were similarly or oppositely regulated across both lines. Potential pathway crosstalk within each cell line was assessed by identifying overlapping DEGs between pathways using GeneOverlap v1.34.0 ^101^, which provides statistical measures of overlap. These results were visualized with bubble plots, and the expression of key overlapping genes was further illustrated using heatmaps.

### WGCNA analysis

The analysis was performed on organoids from the CTL04 line for a total of 66 samples. Protein-coding and long noncoding genes were selected; filtering was then applied to discard not- expressed or low-expressed genes by keeping genes with an expression of at least 40 counts in at least 4 samples. To identify informative genes for network generation, the coefficient of variation (CV) was calculated on log-transformed expression data (vst transformation from DESeq2 package – version 1.38.3) and selection of the genes with the highest CV was applied (35th quantile, 10726 genes). The WGCNA R package (version 1.72-1) was used to generate a signed co-expression network. The correlation matrix was calculated by applying a biweight mid-correlation and then transformed into an adjacency matrix by raising it to the power of β= 14. Topological Overlap Measure was calculated from the adjacency matrix and the relative dissimilarity matrix was used as input for average-linkage hierarchical clustering and gene dendrogram generation. 18 gene modules (plus a grey module grouping not-assigned genes) were defined as branches of the dendrogram using the DynamicTree Cut algorithm (deepSplit=1; minimum cluster size= 60; PAM stage TRUE; cutHeight 0.999; merging modules whose distance is less than 0.15). For module-trait correlation, module eigengens were correlated with exposure conditions dichotomized as a series of categorical variables (dummy variables). To do so, module eigengenes, the first principal component of a given module, were calculated and correlated using Spearman metrics. ME of interesting modules were visualized through exposure conditions to analyze their trends. For module functional analysis, gene ontology enrichment analysis was performed on the genes belonging to each module, with the 10726 genes selected for network generation used as gene universe. The analysis was performed using TopGO as described above; PValue 0.01 and enrichment of 2 were used as significance thresholds.

### Single-cell processing and sequencing

Neural organoids were collected at DIV 50 (+/- 3 days). For each scRNA seq run, 2 exposure conditions were processed, using 3 organoids per condition to obtain technical replicates which were univocally labelled with Cell Multiplexing Oligos (CMOs) (10X Genomics). After rinsing organoids in PBS twice, dissociation was performed following the Stemcell Neural Organoids dissociation protocol ([link](https://www.stemcell.com/how-to-dissociate-3d-neural-organoids-into-single-cell-suspension.html)). Briefly, neural organoids were incubated with 1 ml solution containing 30 U/ml of activated Papain (StemCell) and 125 U/ml of DNaseI (Zymo Research) on orbital shaking conditions (120 rpm, 37 °C, 5% CO2) for 45 minutes. 1 ml of ovomucoid solution (Worthington) was added to stop enzymatic reaction, then digested suspension was pipetted 6 times with a p1000 to break down remaining clumps and centrifuged at 200 RCF for 3 min. Cell pellets were resuspended in cold PBS–0.04% BSA and filtered through 70-μm-pore Flowmi Cell Strainers (Sigma, BAH136800040). Cells were counted and 1 million live cells for each of the replicates were collected and labelled with a unique barcode from the 3’ CellPlex Kit Set A (10X Genomics) according to manufacturer’s instructions ([link](https://cdn.10xgenomics.com/image/upload/v1660261285/support-documents/CG000391_DemonstratedProtocol_CellLabelingwithCellMultiplexingOligo_RevB.pdf)). During each cell counting step, a sample viability >90% was used as a cutoff, excluding all samples with lower viability to prevent CMOs shuffling and thus failure of post-sequencing replicates demultiplexing. After completing the washing steps, 1x 10^5^ cells per replicate were pooled together, subsequently adjusting the cell concentration to be within 1.2-1.6 x 10^6^ cells/ml to obtain an estimated target recovery of 30000 cells/sample. Droplet-based single-cell partitioning and single-cell RNA-Seq libraries were generated using the Chromium Next GEM Single Cell 3’ v3.1 kit (10X Genomics) following manufacturer’s instructions and as detailed in ^102^. Two indexed libraries were equimolarly pooled and sequenced on Illumina NOVAseq 6000 platform using the v3 Kit (Illumina, San Diego, CA) with a customized paired end, dual indexing (26/8/0/98-bp) format according to the recommendation by 10× Genomics. Using proper cluster density, a coverage of around 750 M reads per sample (30000 cells) was obtained for the gene expression, corresponding to at least 25,000 reads/cell whereas for CellPlex Multiplexing Oligos a sequencing depth of 5000 reads/cell was used as recommended by 10X Genomics.

### Data demultiplexing

CMOs demultiplexing was performed through the Cell Rangel Multi pipeline to retrieve the identity of each cell after general QC. The results of Cell Ranger Multi demultiplexing were further validated by comparison with the classification of identities performed by an alternative pipeline, namely deMULTIplex v1.0.2 ^51^. For each run, the sample barcode count matrix of non-empty droplets, as generated by Cell Ranger multi, were given as input to the classification and negative-cell retrival workflow. Furthermore, given that each run contained the same experimental exposure conditions for both lines, a further check on CMOs-based demultiplexing quality was performed by comparing all identities attributed to each line by Cell Ranger Multi and MultiSeq pipeline against the genetic demultiplexing performed through the aggregated call method described by us in ^10^.

### Single-Cell RNA data pre-processing

After the samples demultiplexing only cells expressing more than 100 were retained and each sample was individually submitted to scDblFinder ^103^ for doublets detection and removal. Subsequently, mean absolute deviation (MAD) filtering was applied to the cells each exposure condition individually. MAD-based filters were applied to mitochondrial and ribosomal transcripts content (max. rates: mean rate + (3*MAD)), log transcript and number of detected genes (min. values: mean - (2*MAD)). Additional quality metric flags were added scoring (using tl.score.gene function from) for necroptotic process (GO:0070266), programmed necrotic cell death (GO:0062098), ER (GO:0034976) stress and hypoxia (GO:0001666), the latter metrics were computed condition-wise to avoid the flattening of treatment-driven effects, with only cells passed at least three over the four QCs (score < mean + (3*MAD)). Finally, after integration, dimensionality reduction and leiden-clustering top markers per cluster were used to score and remove cells with high and localized expression of endothelial markers / mesodermal differentiation, ER stress.

### Highly variable genes (HVGs) selection

To maximize the preservation of exposure-driven effects while reducing technical variability, HVGs selection was carried-out with a multi-step approach leveraging the three replicates per condition per PSC line. *i)* Intra condition HVGs: for each cell line and condition highly variable genes were detected using sc.pp.highly_variable_genes function (with options: flavor="seurat", n_top_genes=2000), keeping only genes resulting highly variable in at least *n – 1* replicates. ii) Inter-condition HVGs: computed across conditions per each line. The final HVGs list consisted of of HVGs retrieved in *i)* and common to both PSC upon exposure to the same compound, and inter-exposure HVGs resulting from ii) and common to both PSC lines. In the downstream analyses focused on progenitors and neurons HVGs selection was similarly to *i)* carried out but keeping PSC lines completely separated. Integration via python harmony implementation ^52^ (with default parameters) was carried out solely to coherently, annotate progenitors and neurons populations across the different PSC (Extended Data Fig. 6). After splitting the dataset into neurons and progenitors for in-depth analysis of each cell type, each line was analysed independently, and results were projected on the unintegrated cell types (Fig. 3, Extended Data Fig. 8, [[GitHub link for progenitors](https://giuseppetestalab.github.io/noha/10.2_Hormones_substudy_LineMagnification_Progenitors.html), [GitHub link for neurons](https://giuseppetestalab.github.io/noha/10.2_Hormones_substudy_LineMagnification_Neurons.html)]).

### Differential abundance analysis (DAA)

DAA was carried out using Milo ^53^. for each line independently and oriented to the detection of differences among agonist and antagonist of each compound. Thus, the steps preceding the DAA were set to accordingly and leveraging the technical replicates (see Highly variable genes (HVGs) *i)*). For each macro cell type, condition and PSC line, we extracted spatial neighborhoods showing significant changes in abundance (|logFC| > 1 and FDR < 0.01) and filtered out ambiguous (cells belonging to discordantly enriched neighbors) or sparsely populated neighborhoods (fewer than 20 cells). Barcodes associated with significantly enriched or depleted neighborhoods were marked according to their differentially abundant status (DA) assigning them to corresponding categories ("Enriched", "Depleted", or "NotEnriched"). These labels were stored in the .obs slot of the AnnData object for downstream visualization on the comprehensive UMAP of the corresponding cell type domain.

### WGCNA of differentially abundant (DA) neighbors

To further dissect the transcriptional differences induced by the different compound exposures, we performed weighted gene co-expression network analysis (WGCNA) on DA neighbors and cells. We selected previously marked DA cells from agonist-antagonist comparison, focusing on those categorized as “Enriched” (agonist only) or “Depleted” (antagonist only). For each compound-line combination, conditions with fewer than 100 cells in either DA category were excluded to ensure robustness. Cells were grouped by sample ID and DA status, and pseudobulk expression profiles were generated by aggregating raw counts. Subsequently, the values for each pseudobulk were normalized to the total sum of 1M to accommodate the bulk-oriented pre-processing and filtering. The so-obtained normalized counts stored in an AnnData objects and used for downstream WGCNA via PyWGCNA python package ^104^, with condition and compound metadata annotated accordingly. The gene modules resulting from WGCNA, were filtered according to their ability to separate PSC lines / compound exposures (anova test on module-wise derived PC1 loadings providing as covariate the line, enrichment status combination).

### scRNA-based Differential expression analysis (DEA)

To characterize the transcriptomics differences independently ascribable to progenitors or neuronal changes, after the isolation of the two macro cell types, we generated pseudobulk samples by aggregating cells by replicate (preserving therefore the variability of each exposure-PSC-replicate) to better control false discoveries as suggested in ^105^. Differential expression (DE) analysis was then performed using edgeR ^82^, testing multiple contrasts:

i. Line-shared effects of compound exposure (e.g., agonist vs. DMSO or agonist vs. antagonist), modeled with the PSC line covariate blocked to detect shared responses.
ii. Line-differential effects, evaluating whether the same compound elicits different transcriptional responses across PSC lines.
iii. Per-line effects, assessing compound-induced changes within each line independently, without including cross-line covariates at test time.

In all cases, the design matrix used for filtering, variance estimation, and normalization included the appropriate covariates -when present-reflecting the hypothesis being tested. DE results were filtered at a false discovery rate (FDR) threshold of < 0.05. For per-line analyses, we additionally required an absolute log2 fold change > 2 to retain biologically meaningful differences (All tables with condition-specific DEGs are available at [[GitHub link for progenitors](https://github.com/GiuseppeTestaLab/noha/tree/main/2_scRNASeq/hormones_substudy/hormones_substudy_Progenitors_DEGs), [GitHub link for neurons](https://github.com/GiuseppeTestaLab/noha/tree/main/2_scRNASeq/hormones_substudy/hormones_substudy_Neurons_DEGs)]).

### Image analysis

For the quantifications of the number of nuclei positive to specific markers, we first started by dearraying the TMA and isolating each organoid in each region of the scanner by performing nuclei detection with Stardist ^106^ and fusing overlapping nuclei bounding boxes. For each of the scanned regions, a report highlighting the bounding box for each organoid was produced [[GitHub folder link](https://github.com/GiuseppeTestaLab/noha/tree/main/3_ImageAnalysis/preprocessing/reports)], and for those that overlapped more organoids or yielded any problem, a manual bounding box was produced using the label feature of Napari. Additionally, manual quality checks for each channel were done to evaluate tissue integrity and signal from the stainings [[GitHub link](https://github.com/GiuseppeTestaLab/noha/tree/main/3_ImageAnalysis/preprocessing/tables)]. The final number of organoids used for the analysis is 139.

Nuclei were then recomputed for each single organoid with Stardist and intensity and morphological measurements were quantified with the “regionprops” function from scikit-image ^107^. Morphological parameters measured were ‘area’, ‘bbox’, ‘centroid’, ‘eccentricity’, ‘equivalent_diameter’, ‘extent’, ‘major_axis_length’, ‘minor_axis_length’, ‘orientation’, ‘perimeter’ and ‘solidity’.

For classification of positive nuclei to a specific marker, Otsu’s multi-thresholding method^108^ was applied to the mean intensity of all nuclei in an organoid quantified for the marker channel. Three different thresholds were obtained for each distribution, and the middle one was used as the gate for the definition of positive nuclei in that organoid.

For the quantification of cytoplasmic proteins, Otsu’s multi-thresholding method was used as well on the log-normalized values of the corresponding channels. Similarly to what done for the classification of nuclei, three thresholds were detected and the middle one was used to define positive pixels to the marker. The total number of positive pixels was used to compare the total signal of the cytoplasmic marker across conditions.

In parallel, organoid areas were quantified by combining the nuclei mask with the mask detecting cytoplasmic signal. Nuclei densities were evaluated as the ratio of the number of nuclei to the organoid area.

Differences amongst condition morphological features of all nuclei were analysed through principal component analysis implemented in the scikit-learn package ^109^. The resulting principal components were grouped by organoid for plotting.

For more details, all code is made available at [[GitHub folder link](https://github.com/GiuseppeTestaLab/noha/tree/main/3_ImageAnalysis)]. All the code for image analysis was run with alessiavalenti/imaging:TMA-0.0.1.

## Acknowledgements and funding

We thank present and former members of the Testa lab for collaborating to various extents on this study with both technical help and conceptual discussion. We thank the European School of Molecular Medicine (SEMM) at which D.C., M.T.R., A. V, M.L, G.M., S.S. are/were enrolled as students for their PhD degree program in Systems Medicine. The project, carried out in G.T.’s lab at the European Institute of Oncology (IEO) and at Human Technopole (HT), has been funded by European Union Horizon 2020 research and innovation program grants ENDpoiNTs (825759), NEUROCOV (101057775), RE-MEND (101057604), and by Fondazione Cariplo and Fondazione Telethon joint call (GJC23036). Part of the experimental and analytical work of this study has been carried out at the Tissue processing, Imaging, and Genomics facilities of IEO and HT.

## Author contributions

N.C., and G.T. conceived the project and, with M.T.R., C.E.V., C.C., implemented the experimental and analytical designs; M.T.R. drove the experimental activities with the help of M.L., S.S., B.M., N.C., D.B., and S.T.; G.M., D.C., A.V., C.C., drove the computational work with the help of M.K., and A.T.; G.M., M.T.R., D.C., A.V., M.L., N.C., C.C., S.S., B.M., R.N., L.M., A.M. contributed to analyses, critical discussions and interpretation of the results; B.M. drew figures’ schemes; S.E., P.L., quantified the steroids metabolites in neural organoids; N.C., C.E.V., C.C., and G.T. supervised wet and computational activities; N.C. drafted the paper with input from all authors; N.C. and G.T. finalized the paper. All authors read and approved the submitted paper.

## Data and Code availability

Raw data will be made publicly available upon acceptance of the manuscript for publication.

Full code used for the analyses is available at https://github.com/GiuseppeTestaLab/noha

Processed and annotated data will be made publicly available through relevant portals, specific links indicated in the GitHub readme file.

### Report Cards and Mining

Acknowledging the wealth and multi-scale nature of the data produced in this study, which integrates bulk and single-cell transcriptomics, high-throughput imaging, and targeted steroidomics, we have developed a public resource to ensure maximum accessibility and utility for the scientific community.

For each of the seven hormonal pathways investigated, we provide a dedicated "report card" containing a schematic summary of the main observations drawn by our analyses. This allows fellow researchers to gain a rapid, high-level understanding of the key findings. A more comprehensive asset complements this initial overview: a mining of the data organized in this public repository serving as a portal for deep data exploration, offering detailed descriptions of our results and, crucially, contextualizing them within the current body of scientific knowledge. This transforms our atlas from a static publication into a dynamic resource designed to be actively used, explored, and updated. While we have meticulously curated this initial release, we envision it as a living atlas that will evolve and improve in the future. This was elaborated as a collective effort distributed across scientists with different backgrounds and seniority. Consequently, even if the same structure with thematic sub-chapters was elaborated for each hormonal pathway, the level of details included for each pathway is still heterogeneous, mainly because the amount of available public literature is very different across pathways. We plan to continuously refine the harmonisation of the relevant datasets and information on the portal, hoping that the scientific community will further contribute to it as a collective exercise, aligned to the principles and practice of the Human Cell Atlas. Our vision is that this resource will foster further discoveries and provide a framework for investigating the impact of endocrine signalling in human neurodevelopment, and more broadly, the developmental origins of neuropsychiatric traits.

If you notice any typos or glitches, feel welcome to contact us to contribute and improve this resource.

The work is accessible here [[link](https://github.com/GiuseppeTestaLab/noha/blob/main/1_bulkRNASeq/7_DataMining/0.Index.md)].

## Declaration of generative AI and AI-assisted technologies in the writing process

During the preparation of this work the authors used AI tools in order to improve language and readability. After using these tools, the authors reviewed and edited the content as needed and take full responsibility for the content of the publication.

